# Co-lethality of *LEU2* and *BAP2* is carbon source dependent and is overcome by *BUL1* deletion in *Saccharomyces cerevisiae*

**DOI:** 10.1101/2024.10.05.616764

**Authors:** S Maheswaran, Paike Jayadeva Bhat

**Affiliations:** Department of Biosciences and Bioengineering, Indian Institute of Technology, Bombay. Mumbai. India

**Keywords:** Leucine permease, Arrestin related trafficking adaptor, Post translational regulation, Leucyl tRNA synthetase, Target of Rapamycin and Chronological life span

## Abstract

In a leucine requiring *Saccharomyces cerevisiae* strain, deletion of *BAP2*, a high affinity leucine permease confers lethality in presence of fermentable sugars, but not other carbon sources. Deletion of *BUL1*, an adaptor overcomes this lethality. *BUL1* deletion augments leucine import through accumulation of Bap3 and Bap2 in the plasma membrane and reduces their turnover. We suspect *BUL1* as a cognate adaptor for ubiquitylation of these permeases.

## Introduction

From bacteria to humans, amino acids are either self-synthesized or outsourced. *Saccharomyces cerevisiae* synthesizes all of the proteinogenic amino acids from inorganic salts like Ammonium sulfate and organic carbon source glucose (Godard et al., 2007). Additionally, it has a battery of broad or narrow specific permeases to import amino acids from the external milieu (Zhang et al., 2018; Bianchi et al., 2019).

Amino acid permeases are regulated at transcriptional and post translational levels. e.g. Transcription of Put4, a proline permease and Gap1, the general amino acid permease is upregulated when *Saccharomyces cerevisiae* thrives on a poor-quality nitrogen and downregulated in a good-quality nitrogen (Jauniaux et al., 1987; Jauniaux and Grenson, 1990). When a poor-quality nitrogen is replaced by a good-quality nitrogen, Gap1 in plasma membrane are ubiquitylated, endocytosed and degraded in vacuole (De Craene et al., 2001). The last case makes an example of post translational regulation.

Permease ubiquitylation is achieved by a duo of ubiquitin ligase and adaptor. Rsp5, the ubiquitin ligase in association with adaptors ubiquitylate permeases (Lin et al., 2008). The adaptor family includes membrane bound adaptors such as Ear1, Ssh4, Tre1, Tre2, Bsd2 and 14 soluble, cytoplasmic adaptors such as Art1/Ldb19, Art2/Ecm21, Art3/Aly2, Art4/Rod1, Art5, Art6/Aly1, Art7/Rog3, Art8/Csr2, Art9/Rim8, Art10, Bul1, Bul2, Bul3 and Spo23 (Hettema et al., 2004; Stimpson et al., 2006; Leon et al., 2008; Lin et al., 2008; Sardana and Emr, 2021; Zbieralski and Wawrzycka, 2022). The soluble ones are called as arrestins or Arrestin Related Trafficking adaptors, shortly ARTs.

Factors such as presence, absence and concentration of substrates, stress like high pressure, antibiotics, heavy metals, shift in carbon, nitrogen sources and various other signals drive permease downregulation through ubiquitylation (Lai et al., 1995; Horak and Wolf, 1997; Krampe et al., 1998; Beck et al., 1999; Abe and Iida, 2003; Liu and Culotta, 1999; Gitan and Eide, 2000; Blondel et al., 2004; Felice et al, 2005; Paiva et al., 2009; Nikko et al., 2008; Hatakeyama et al., 2010; Becuwe et al., 2012; Zhao et al., 2013; Suzuki et al., 2013; Fernandez-Murray et al., 2013; Crapeau et al., 2014; Talaia et al., 2017; Fujita et al., 2018; Hovsepian et al., 2017; Hovsepian et al., 2018; Savocco et al., 2019; Tanahashi et al., 2021; Megarioti et al., 2021; Kozu et al., 2021; Robinson et al., 2022).

Usually, an adaptor directs Rsp5 ligase ubiquitylate a set of permeases and a permease has more than one adaptor for its ubiquitylation (Nikko and Pelham, 2009; O’Donnell and Schmidt, 2019; Kahlhofer et al., 2021; Zbieralski and Wawrzycka, 2022).

In this study, we assessed the dispensability of a few prominent amino acid permeases of *Saccharomyces cerevisiae* of s288c background in synthetic media. We found that deletion of *BAP2*, a high affinity leucine permease resulting in growth cessation or lethality. This lethality was overcome by deletion of *BUL1*, which resulted in accumulation of Bap3 in the plasma membrane. Our studies imply Bul1 as a cognate adaptor of Bap2 and Bap3.

## Results

### LEU2 and BAP2 are co-lethal

Deletion of *LEU2*, an enzyme in leucine biosynthesis is lethal for *Saccharomyces cerevisiae* strains growing in synthetic medium (Cohen and Engelberg, 2007). *LEU2* complementation or expression of leucine permeases *BAP2*, *TAT1* in a multicopy plasmid overcomes this lethality. Also, deletion of *BAP2* and *LEU2* manifests growth retardation or lethality (Nigavekar and Cannon, 2002; Usami et al., 2014). It is assumed that reduced leucine import due to the fierce competition from other amino acids in excess in the media is the cause for lethality (Usami et al., 2014).

We wished to know if *LEU2* establishes co-lethality with other major amino acid transporters like *BAP3*, *TAT1*, *AGP1*, *GNP1* which transport leucine and are induced by it (Didion et al., 1998; Regenberg et al., 1999). We have also included *TAT2* in this study. Strains lacking *LEU2* and transporters were studied for growth in SC Glucose and SC Galactose (Fig. 1A).

**Figure 1.**
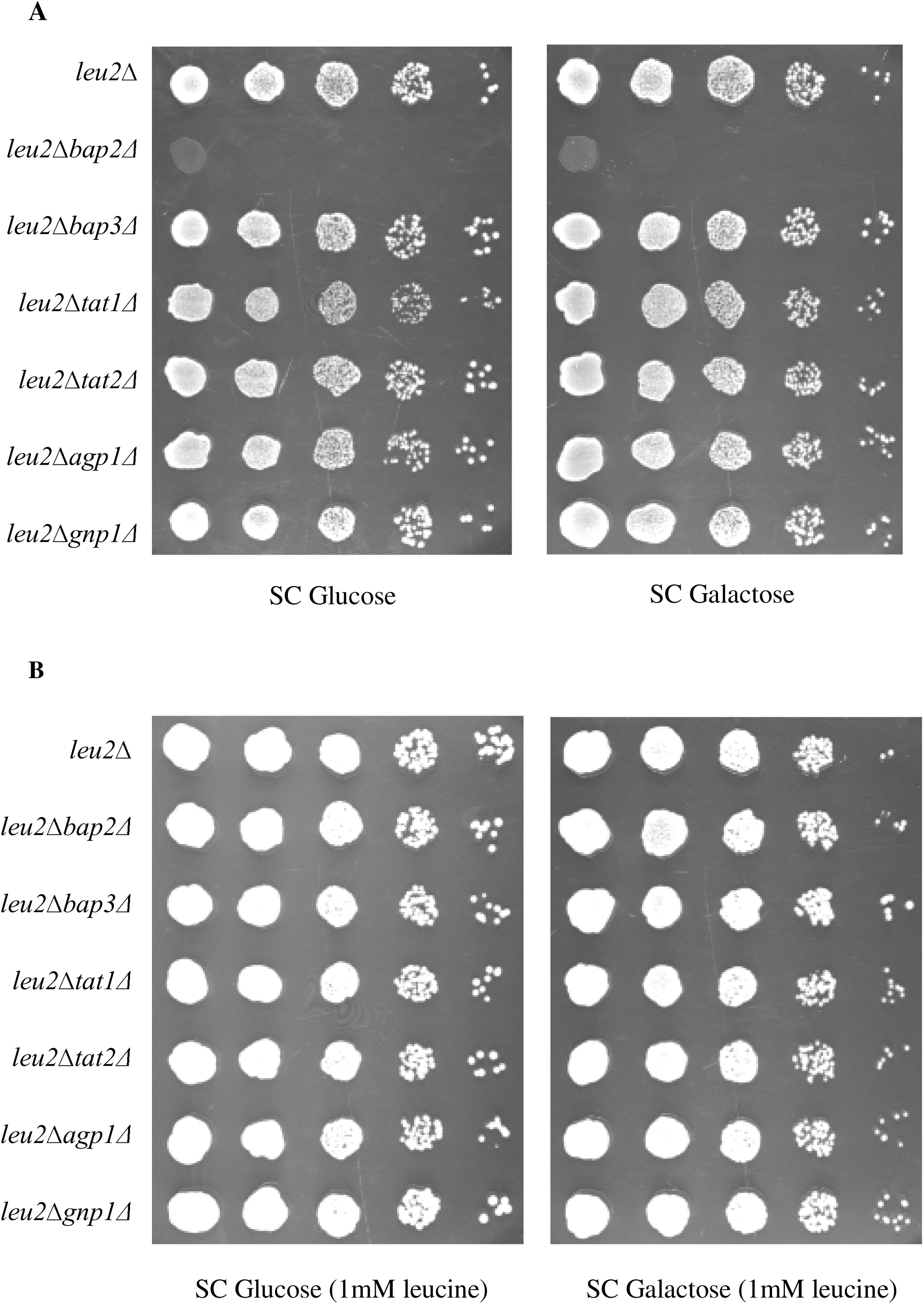
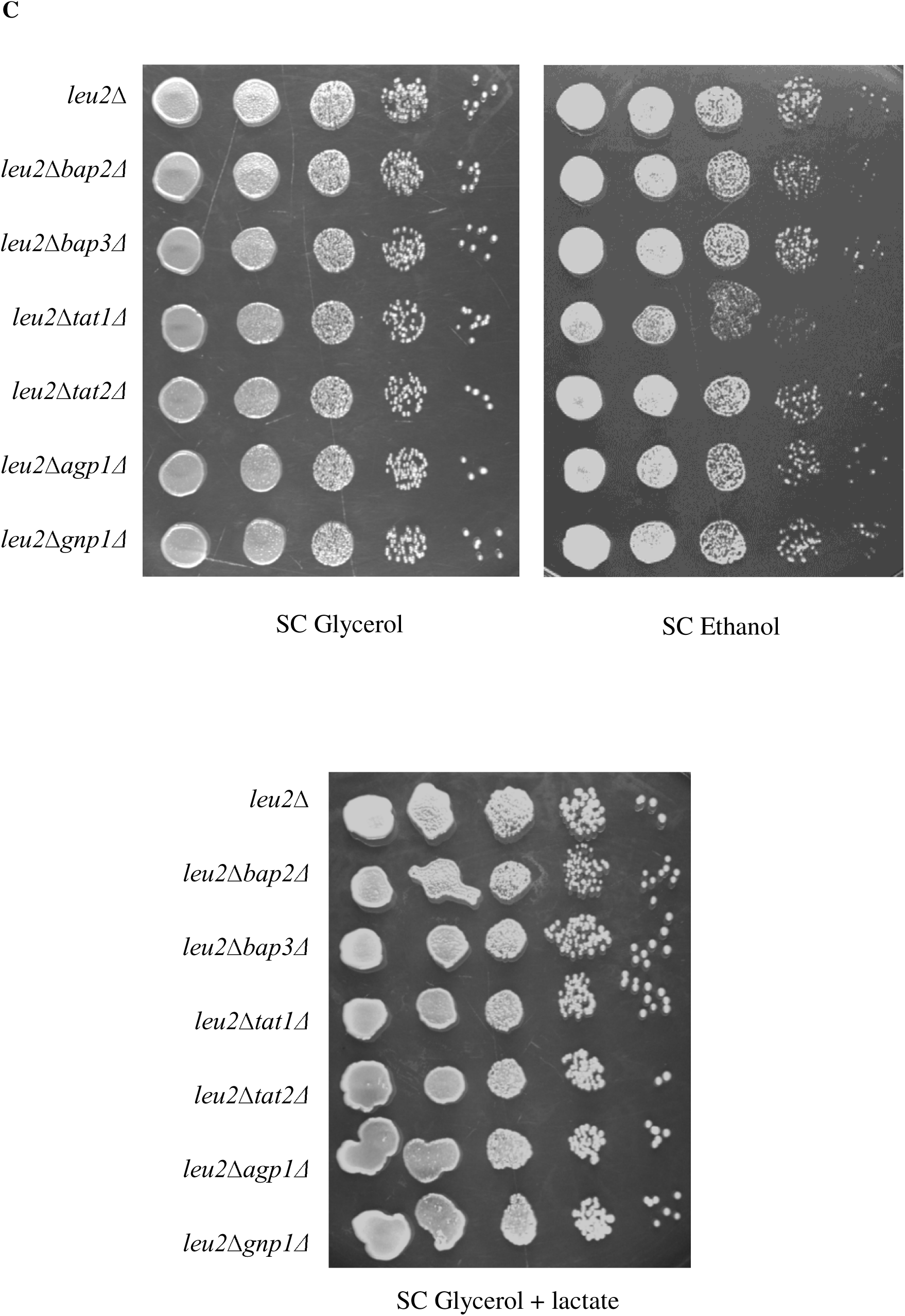
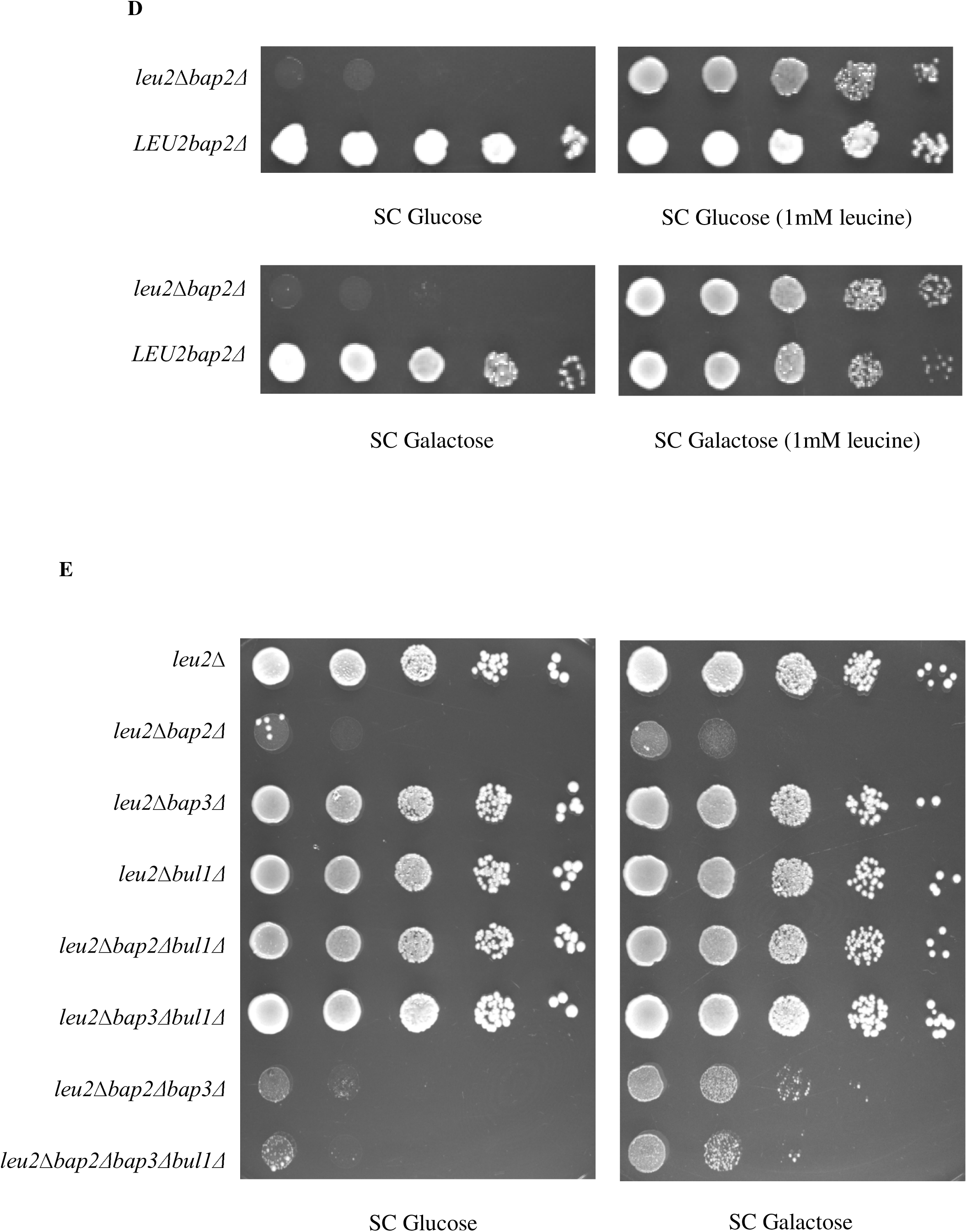
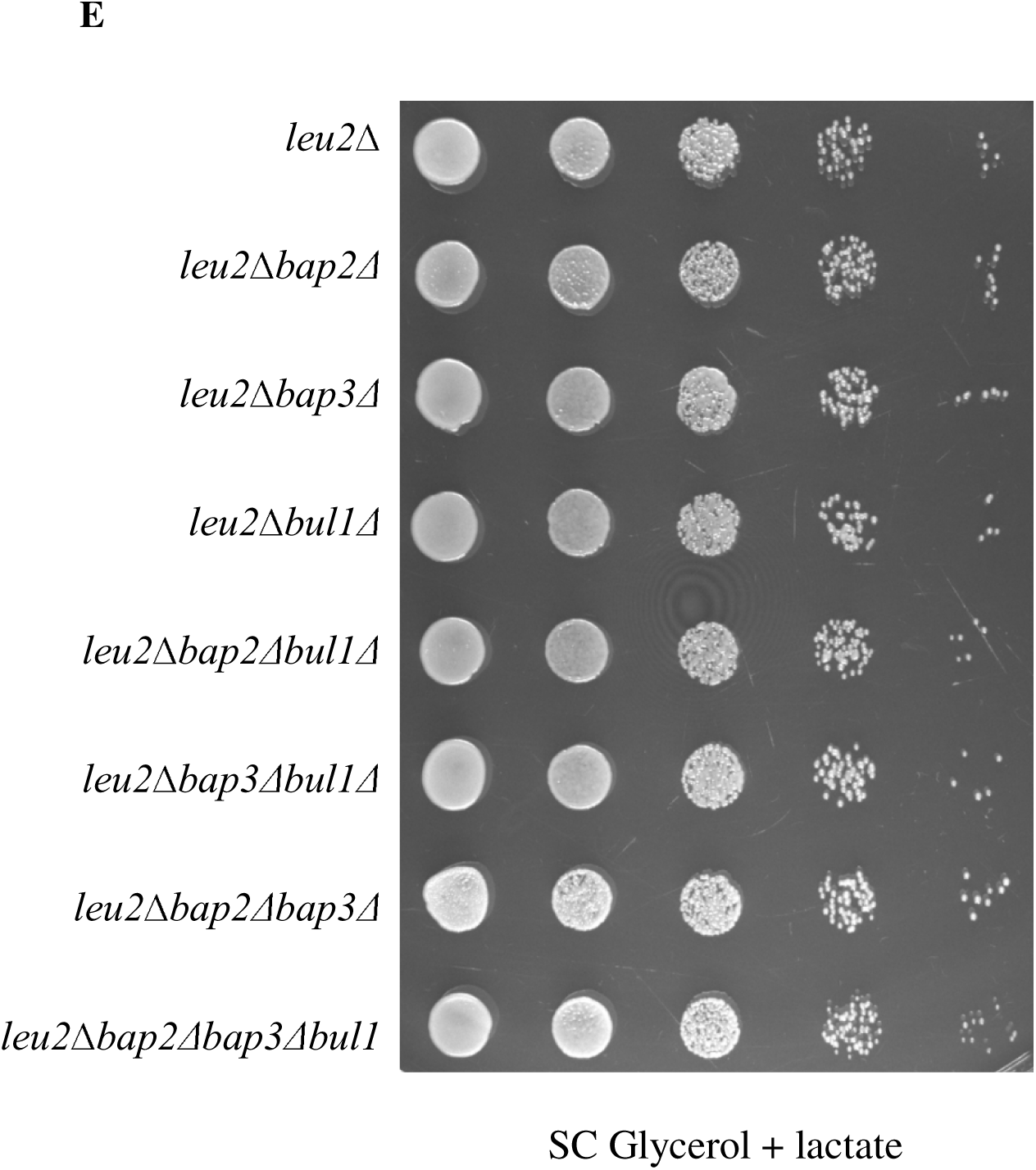
Deletion of *BAP2* and *LEU2* confer co-lethality in a carbon source dependent manner. (A) *leu2*Δ, *leu2*Δ*bap2*Δ*, leu2*Δ*bap3*Δ*, leu2*Δ*tat1*Δ*, leu2*Δ*tat2*Δ*, leu2*Δ*agp1*Δ and *leu2*Δ*gnp1*Δ strains grown in SC Glycerol + lactate broth were spotted on SC Glucose and SC Galactose. (B) *leu2*Δ, *leu2*Δ*bap2*Δ*, leu2*Δ*bap3*Δ*, leu2*Δ*tat1*Δ*, leu2*Δ*tat2*Δ*, leu2*Δ*agp1*Δ and *leu2*Δ*gnp1*Δ strains grown in SC Glycerol + lactate broth were spotted on SC Glucose (1mM leucine) and SC Galactose (1mM leucine). (C) *leu2*Δ, *leu2*Δ*bap2*Δ*, leu2*Δ*bap3*Δ*, leu2*Δ*tat1*Δ*, leu2*Δ*tat2*Δ*, leu2*Δ*agp1*Δ and *leu2*Δ*gnp1*Δ strains grown in SC Glycerol + lactate broth were spotted on SC Glycerol, SC Ethanol and SC Glycerol + lactate. (D) *leu2*Δ*bap2*Δ and *LEU2bap2*Δ strains grown in SC Glycerol + lactate broth were spotted on SC Glucose, SC Glucose (1mM leucine), SC Galactose and SC Galactose (1mM leucine). (E) *leu2*Δ, *leu2*Δ*bap2*Δ*, leu2*Δ*bap3*Δ*, leu2*Δ*bul1*Δ*, leu2*Δ*bap2*Δ*bul1*Δ*, leu2*Δ*bap3*Δ*bul1*Δ*, leu2*Δ*bap2*Δ*bap3*Δ and *leu2*Δ*bap2*Δ*bap3*Δ*bul1*Δ strains grown in SC Glycerol + lactate broth were spotted on SC Glucose, SC Galactose and SC Glycerol + lactate. All images were captured after incubating the petri plates for 2-3 days at 30°C.

Only the *leu2*Δ*bap2*Δ strain manifested the lethality. We think that devoid of Bap2, the high affinity leucine permease, leucine is not imported in adequacy to sustain growth. Other permeases possibly face difficulty in leucine import due to better affinity or/and abundance of other amino acids in SC Glucose/Galactose media (Regenberg et al., 1999; Bianchi et al., 2019; Usami et al., 2014). The lethality disappears when the leucine in the medium is increased 5 times (1mM) the usual concentration (0.2mM) (Fig. 1B).

Interestingly, the lethality also disappeared when the sugars were replaced by ethanol or glycerol or glycerol plus lactate etc. (Fig. 1C). Nevertheless, the *LEU2bap2*Δ strain grew well in SC Glucose and SC Galactose (Fig. 1D).

*trp1*Δ*erg6*Δ strain is a tryptophan requiring cum import impaired mutant which suffers lethality in synthetic medium (Umebayashi and Nakano, 2003). Tryptophan increment or deletion of *BUL1* restored growth, by accumulating the tryptophan permease Tat2 in the plasma membrane. Bul1 is an adaptor which assists Rsp5 ubiquitin ligase in permease ubiquitylation (Yashiroda et al., 1998; Merhi and Andre, 2012; O’Donnell, 2012).

In this case too *BUL1* deletion in the *leu2*Δ*bap2*Δ strain restored growth in SC Glucose and SC Galactose (Fig. 1E). The growth of *leu2*Δ*bap2*Δ*bul1*Δ strain was dependent on *BAP3*, but not other permeases like *TAT2*, *GNP1* or *AGP1* (Fig. S1A,B). We suspected Bap3 accumulation in the plasma membrane of *leu2*Δ*bap2*Δ*bul1*Δ strain and tracked its localization.

### BUL1 deletion augments leucine import

*leu2*Δ*bap2*Δ and *leu2*Δ*bap2*Δ*bul1*Δ strains were made *LEU2* positive, enabling them to grow in SC Glucose and SC Galactose media, for microscopy and western blotting studies.

*LEU2bap2*Δ*bul1*Δ strain had enriched Bap3*-*GFP distribution in the cell’s periphery than *LEU2bap2*Δ strain (Fig. 2A,B). Such pattern was observed between *leu2*Δ*bul1*Δ and *leu2*Δ strains also (Fig. 2C,D). The GFP signal was fainter in SC Galactose. We supposed the peripheral Bap3-GFP as the plasma membranous Bap3-GFP.

**Figure 2.**
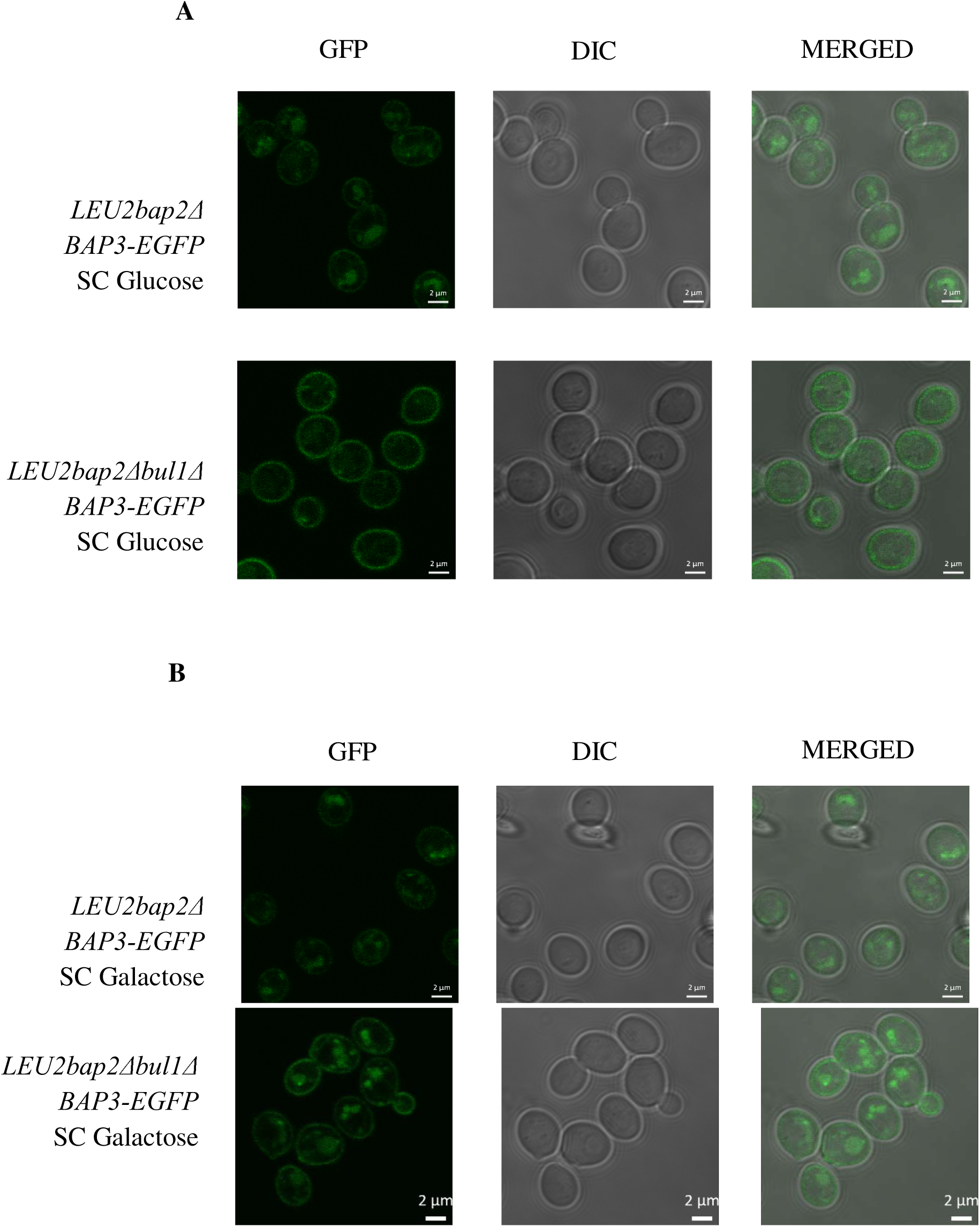
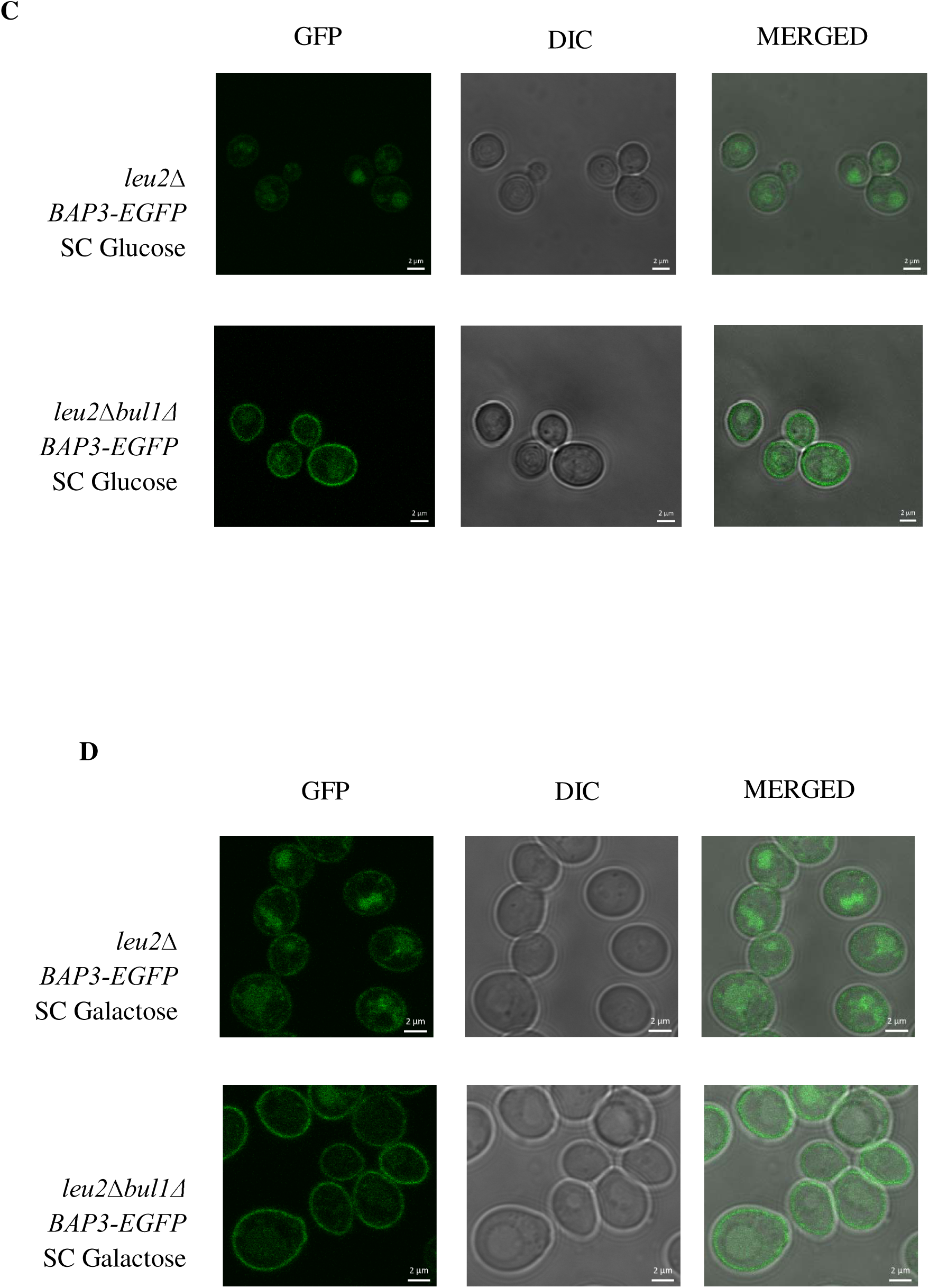
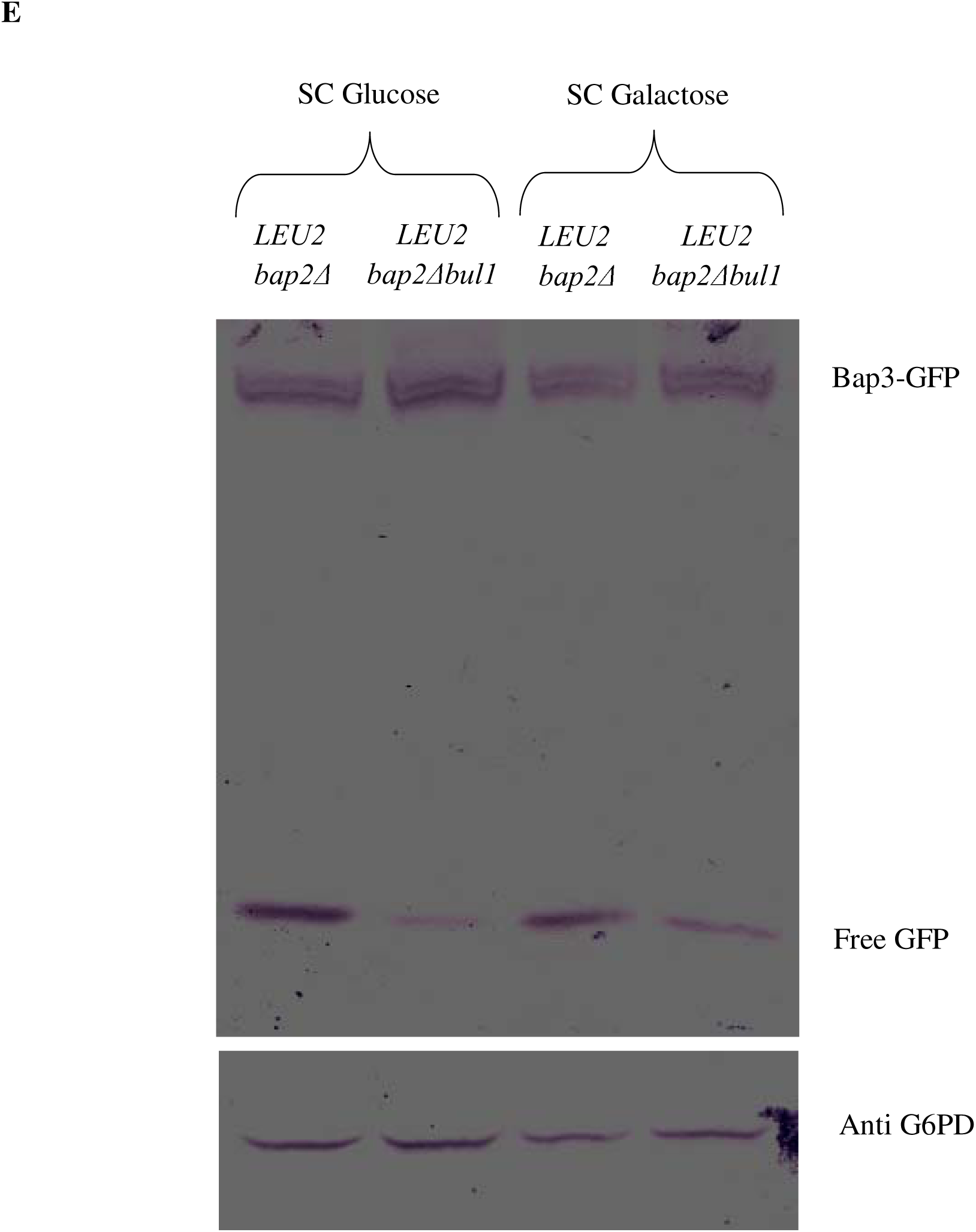
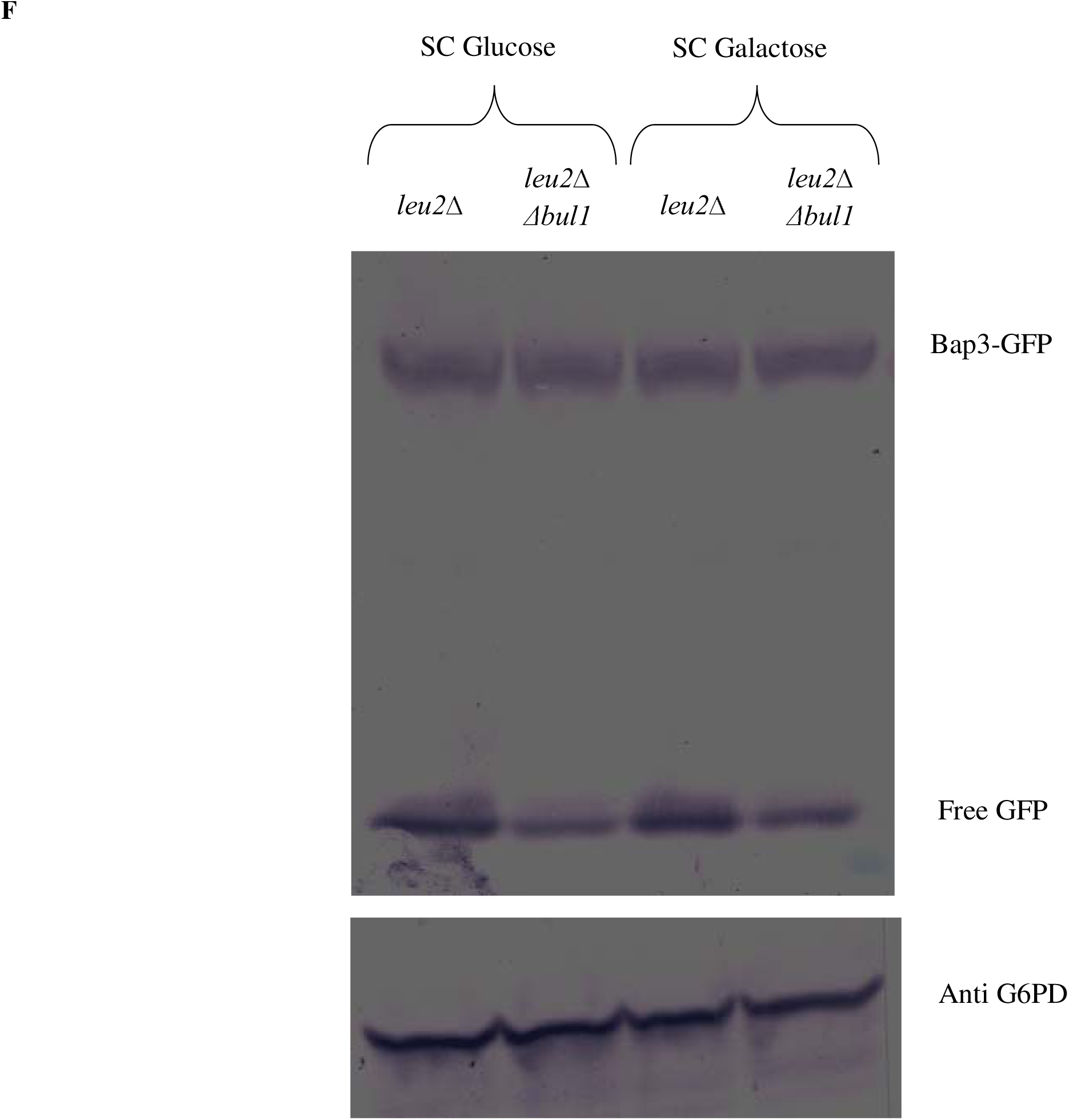
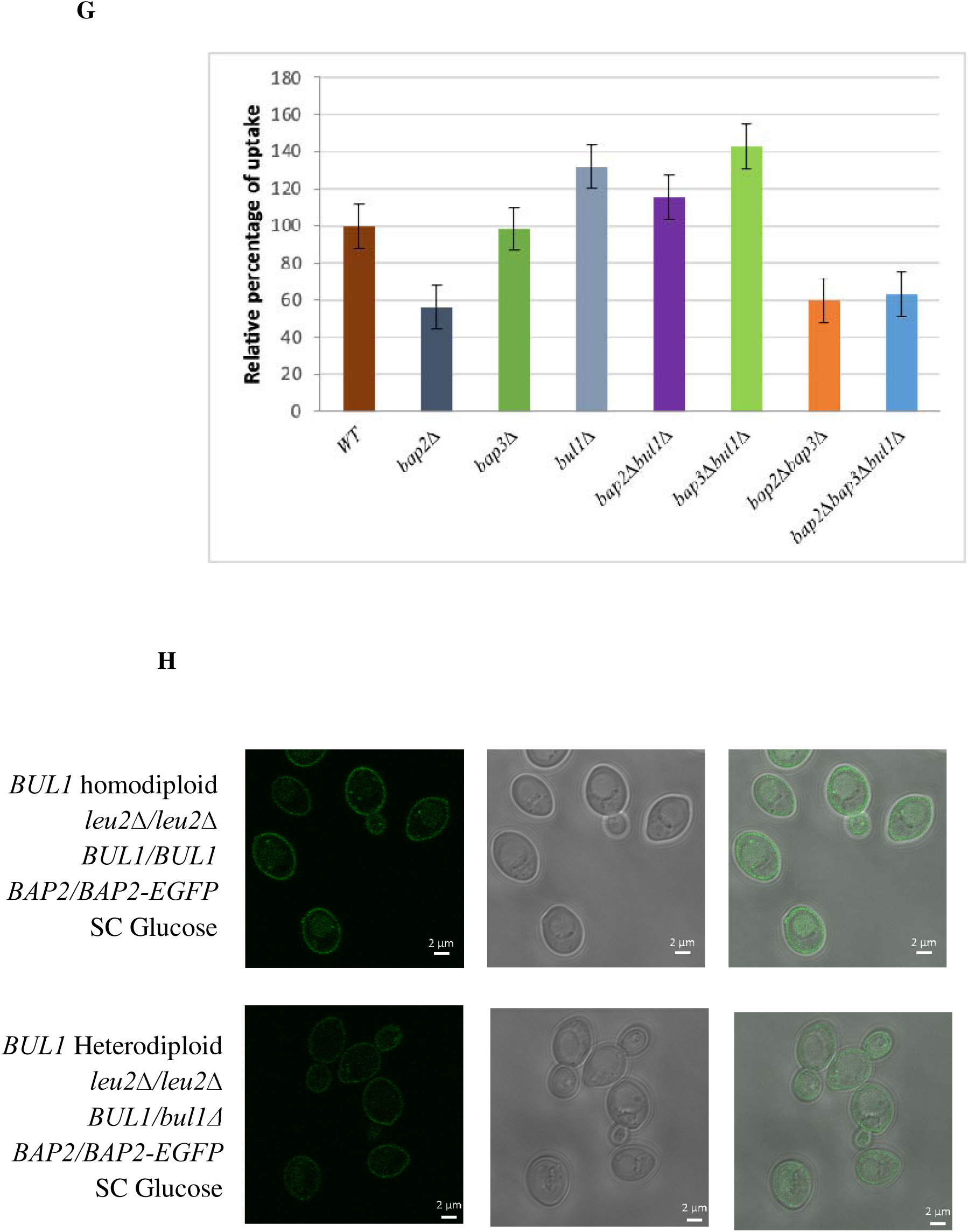
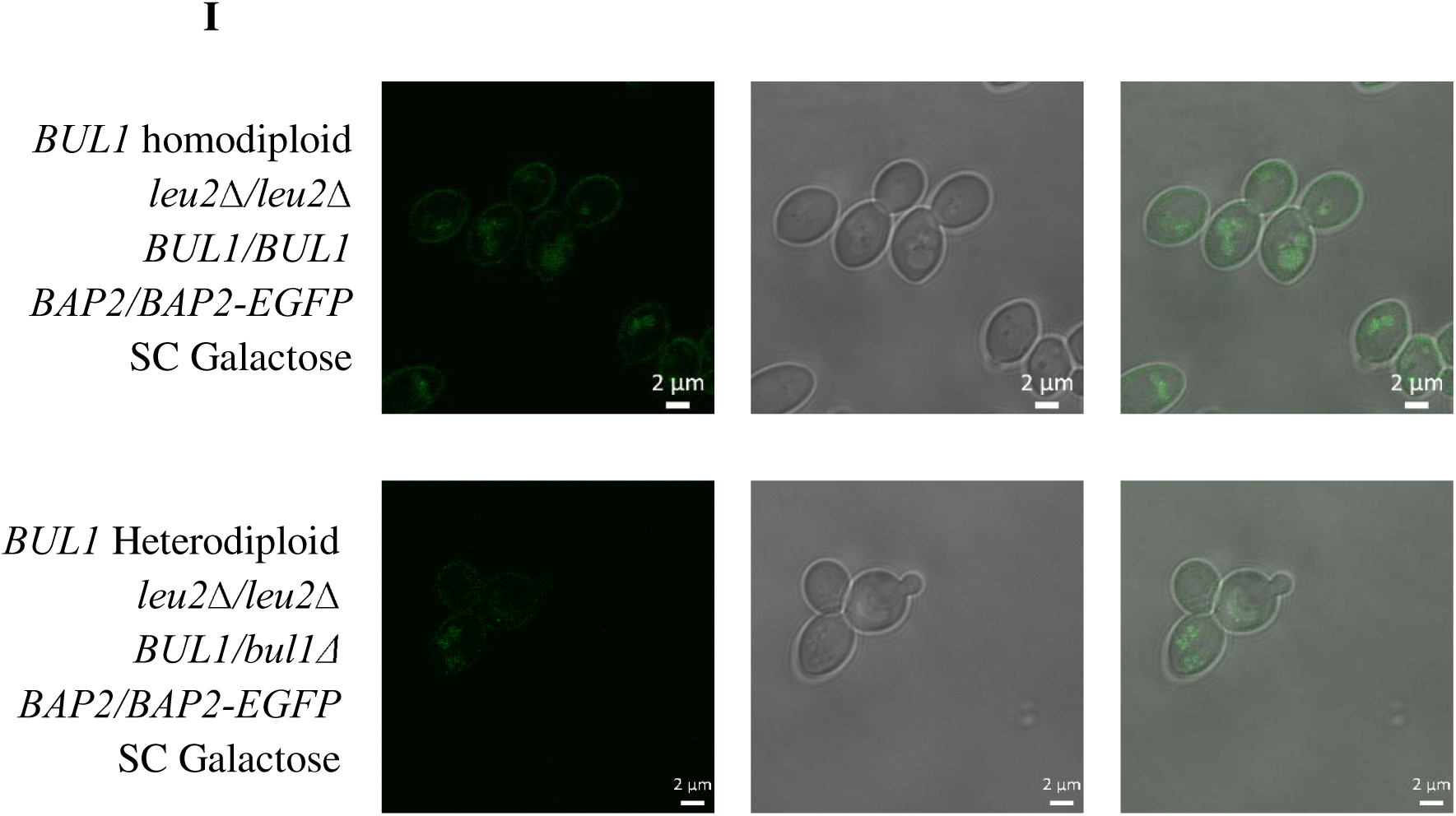
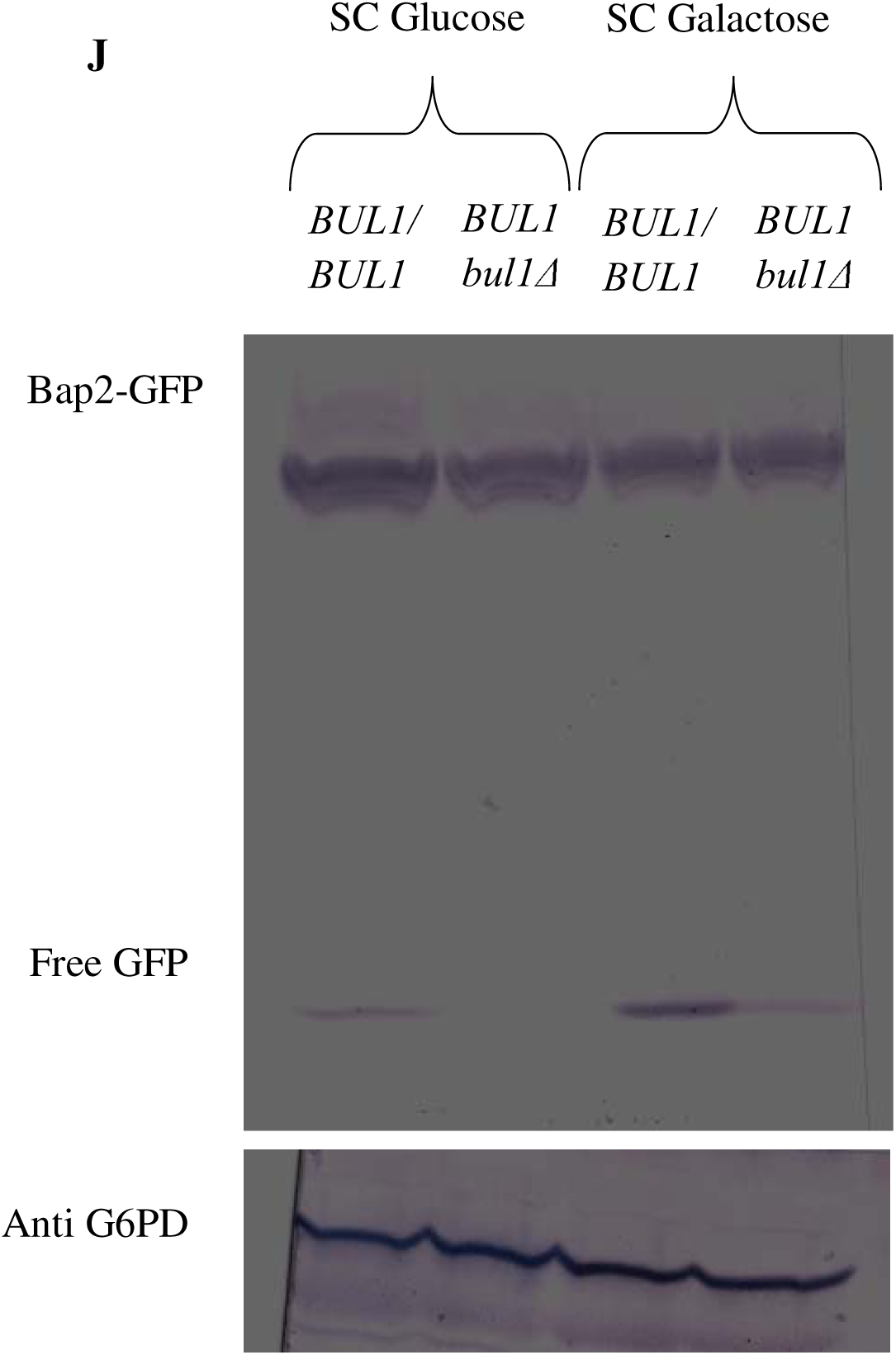
*BUL1* deletion leads to accumulation of Bap3-GFP, Bap2-GFP in the plasma membrane and reduces their turnover. (A) *LEU2bap2*Δ *BAP3-GFP* and *LEU2bap2*Δ*bul1*Δ *BAP3-GFP* strains grown in SC Glycerol + lactate broth were sub-cultured in SC Glucose up to mid logarithmic phase, viewed under microscope. (B) Like 2A, sub-cultured in SC Galactose. All panels for microscopic images read Scale-2µM. (C) *leu2*Δ *BAP3-GFP* and *leu2*Δ*bul1*Δ *BAP3-GFP* strains grown in SC Glycerol + lactate broth were sub-cultured in SC Glucose up to mid logarithmic phase, viewed under microscope. (D) Like 2C, sub-cultured in SC Galactose. All panels for microscopic images read Scale-2µM. (E) Immunoblotting of cell lysates of *LEU2bap2*Δ *BAP3-GFP* and *LEU2bap2*Δ*bul1*Δ *BAP3-GFP* strains, grown as mentioned in 2A and 2B. Anti-GFP and anti-G6PD antibodies (loading control) were used. Free GFP emanates from vacuolar degradation of Bap3-EGFP (F) Immunoblotting of total cell lysates of *leu2*Δ and *leu2*Δ*bul1*Δ *BAP3-GFP* strains, grown as mentioned in 2C and 2D. Anti-GFP and anti-G6PD antibodies (loading control) were used. (G) *LEU2* transformants of strains used in Fig. 1E were assayed for import of C14-leucine in SC Glucose. (H) *BUL1* homodiploid (*leu2*Δ*/leu2*Δ, *BUL1*/*BUL1*, *BAP2*/*BAP2*-GFP) and *BUL1* heterodiploid (*leu2*Δ*/leu2*Δ, *BUL1*/*bul1*Δ, *BAP2*/*BAP2*-GFP) strains grown in SC Glycerol + lactate broth were sub-cultured in SC Glucose up to mid logarithmic phase, viewed under microscope. (I) Like 2G, sub-cultured in SC Galactose. All panels for microscopic images read Scale-2µM. (J) Immunoblotting of cell lysates of *BUL1* homodiploid (*leu2*Δ*/leu2*Δ, *BUL1*/*BUL1*, *BAP2*/*BAP2*-GFP) and *BUL1* heterodiploid (*leu2*Δ*/leu2*Δ, *BUL1*/*bul1*Δ, *BAP2*/*BAP2*-GFP) strains grown as mentioned in 2H and 2I. Anti-GFP and anti-G6PD antibodies (loading control) were used. Free GFP emanates from vacuolar degradation of Bap2-EGFP.

The cell lysates of *LEU2bap2*Δ*bul1*Δ and *leu2*Δ*bul1*Δ strains had less free GFP than *LEU2bap2*Δ and *leu2*Δ strains respectively (Fig. 2E,F). Free GFP comes from vacuole aftermath the degradation of Bap3-GFP, as GFP is resistant for vacuolar degradation (Nikko and Pelham, 2009). However, we observed little or no change in the full length Bap3-GFP between strains.

These findings suggested that *BUL1* deletion leads to accumulation of Bap3 in the plasma membrane and reduces vacuolar Bap3 degradation. Perhaps it is the Bap3 accrued in the plasma membrane of *leu2bap2*Δ*bul1*Δ which imports enough leucine to overcome the lethality.

To verify Bap3 accumulation in plasma membrane, C14-Leucine import was studied (Fig. 2G). To enable grow in SC Glucose, all strains were made leucine prototrophs. In this accord, strains are mentioned in the following manner. e.g. *leu2*Δ as WT, *leu2*Δ*bap2*Δ as *bap2*Δ, *leu2*Δ*bap3*Δ as *bap3*Δ and so forth.

C14-Leucine import by WT was regarded as 100%. Deletion of *BAP2* and *BAP3* led to 40% and 10% dip respectively and agree with earlier reports (Grauslund et al., 1995; Didion et al., 1998). This revealed *BAP2* as a prominent and *BAP3* as a minor leucine permease in SC Glucose. Deletion of *BAP2* and *BAP3* together caused 40% dip.

*BUL1* deletion in a *bap2*Δ strain increased the import which was solely reliant on Bap3. The increment was substantial as it was from 40% drop to 15% in excess than WT. This supports our notion that the Bap3-GFP enrichment at the cell periphery is indeed in the plasma membrane.

*BUL1* deletion in the *bap3*Δ strain also improved the import, which diminished on *BAP2* deletion. Perhaps *BUL1* deletion, also accumulates Bap2 in the plasma membrane. We think that the enhanced import of *bul1*Δ strain is due to concurrent accumulation of Bap2 and Bap3.

We bid to know if Bap2 accumulates in the plasma membrane due to *BUL1* deletion. Whereas, tagging of GFP at *BAP2’s* C terminal in *leu2*Δ*bap3*Δ and *leu2*Δ strains was possible, attempts in *leu2*Δ*bap3*Δ*bul1*Δ and *leu2*Δ*bul1*Δ strains were futile.

Bap2 localization was studied in diploids. Cautiously, in each strain we tagged one copy of *BAP2* and deleted one copy of *BUL1*. The Bap2-GFP localization was indifferent between *BUL1* homodiploid and *BUL1* heterodiploid (Fig. 2H,I). Both strains had prominent peripheral and less internal Bap2-GFP localization. Bap2-GFP signal was fainter in SC Galactose.

C terminal of Bap2 is required for its basal turnover and its truncation increases leucine import (Grauslund et al., 1995; Omura et al., 2001). Modification of Bap2 C terminal leads to enhanced growth in a leucine limiting medium (Wolf et al., 2014). In this accord, we think GFP tagging has altered the typical turnover pattern of Bap2 resulting in indifferent or marginally different localization pattern between the strains.

Yet, the free GFP was lesser in the *BUL1* heterodiploid than *BUL1* homodiploid (Fig. 2J), suggesting a possible reduced endocytic turnover of Bap2.

## Discussion

*Saccharomyces cerevisiae* has many plasma membrane bound amino acid permeases with overlapping substrate specificity. These permeases are not expressed concomitantly. This implies a division of labor where a prominent permease in a nutrient niche would become dispensable or less important in another niche, where another permease plays the prominent role.

A way to decipher this essentiality or dispensability of permeases is to study the growth of permease knockout strains in varying nutrient media. Deletion of broad specificity permease *GNP1* but not its paralog *AGP1*, severely impairs serine uptake (Esch et al., 2020).

In this study, we used a *leu2*Δ strain and establish *BAP2* as a prominent leucine permease in SC Glucose or Galactose medium. The co-lethality of *LEU2* and *BAP2* is already a reported one (Nigavekar and Cannon, 2002; Usami et al., 2014) and we show that this manifests in presence of fermentable sugars but not in non-fermentable carbon sources.

Galactose and glycerol augment manifold amino acid import, particularly leucine than glucose (Peter et al., 2006). Though Leucine import is manifold in galactose, deletion of *BAP2*, the major contributor in the surge possibly leads to a situation like glucose, causing lethality. In cases of Glycerol and ethanol, the surge may be due to upregulation of permease(s) other than *BAP2*. *BAP2* is indispensable for a *leu2*Δ strain in SC Glucose or SC Galactose, yet dispensable in SC Glycerol or SC Ethanol.

*BUL1* deletion overcomes the lethality, requiring *BAP3*. Increased peripheral Bap3 and enhanced C14-Leucine import suggest Bap3 accumulation in the plasma membrane due to *BUL1* deletion. Going by Bul1’s canonical role, it is possible that Bul1 is a cognate adaptor for Bap3 ubiquitylation, a driving factor permease turnover.

We hypothesize that in absence of Bul1, Bap3 is not aptly ubiquitylated and accumulates in plasma membrane. Substantiating our notion, the free GFP population is remarkably reduced in strains deleted for *BUL1*. This surplus Bap3 population in the plasma membrane of *leu2*Δ*bap2*Δ*bul1*Δ strain augments the leucine import to overcome the co-lethality of *LEU2* and *BAP2*. We think that there may exist ways other than impeded Bap3 ubiquitylation to explain the observations.

We have learnt that BY4741 strain used in this study has a loss-of-function *BUL2* allele (*BUL2^F883L^*) (Kwan et al., 2011). We have confirmed the existence of this variation. From the standpoint of reports where Bul1 and Bul2 share several common permeases to ubiquitylate (O’Donnell and Schmidt, 2019; Kahlhofer et al., 2021), possibly the suppression of lethality is due to deletion and loss of function of *BUL1* and *BUL2* respectively. To verify we have expressed a functional *BUL2* (from W303-1A strain) in *leu2*Δ*bap2*Δ*bul1*Δ strain and found no deviation in growth. But, the lethality resurfaced when *BUL1* was expressed (Fig. S2).

We do not rule out the possibility that Bap3 is ubiquitylated by Bul2. Yet we limit our inference to say that deletion of *BUL1* is sufficient to overcome the lethality.

The workability of the plasmid borne *BUL2* was verified by rescuing the heat sensitive phenotype of the *bul1*Δ*bul2*Δ strain (here *bul1*Δ*BUL2^F883L^*) in Synthetic Minimal Tyrosine Glucose medium at 37°C (Fig. S3).

*BUL2* deletion reduces chronological lifespan (CLS) and prolongs replicative lifespan (RLS), due to a suspected increase in TORC1 (Target of Rapamycin Complex 1) activity, as a result of enriched intracellular amino acid pool (Kwan et al., 2011). In another study, CLS of a *leu2*Δ strain was found extended by leucine prototrophy or increased leucine concentration in the medium (Alvers et al., 2009).

Leucine is the most frequent amino acid encoded by the genes (Echols et al., 2002) and Leucyl tRNA synthetase, Cdc60 is the most abundant of its kind (Ghaemmaghami et al., 2003). Cdc60 (yeast) and LARS (Leucyl tRNA synthetase in mammals) serve as intracellular leucine sensors. In presence of leucine, they activate TORC1 (yeast) and mTORC1 (mammals) to promote translation and cell proliferation (Bonfils et al., 2012, Han et al., 2012, Segev and Hay, 2012). From this stance, we wonder whether TORC1 activation is higher in *leu2*Δ*bul1*Δ cells as they import 30% more leucine than *leu2*Δ cells? If yes, how would it affect CLS and RLS?

Strains *leu2*Δ*bap2*Δ*, leu2*Δ*bap2*Δ*bap3*Δ and *leu2*Δ*bap2*Δ*bap3*Δ*bul1*Δ experiencing ∼40% dip in leucine import (Fig. 2G) are also the ones to suffer lethality in SC Glucose (Fig. 1E). We think that the dip establishes a sub-optimal intracellular leucine environment, which may lead to lethality in two possible ways, (1) Cdc60 is unable to activate TORC1 efficiently. Inactive or poorly active TORC1 does not promote translation or proliferation leading to growth cessation.

(2) Cdc60 is unable to charge the tRNA^Leu^, leading to stalling of translation, subsequently growth.

It is anti-intuitive that a leucine concentration at which Cdc60 activates TORC1, is below the Km of Cdc60 to charge tRNA^Leu^. In this accord we favor the former possibility. If the predictions are true, a Cdc60 mutant that activates TORC1 or charge tRNA^Leu^ in sub-optimal intracellular leucine concentration should suppress the lethality.

## Supplementary figures

**Figure S1.**
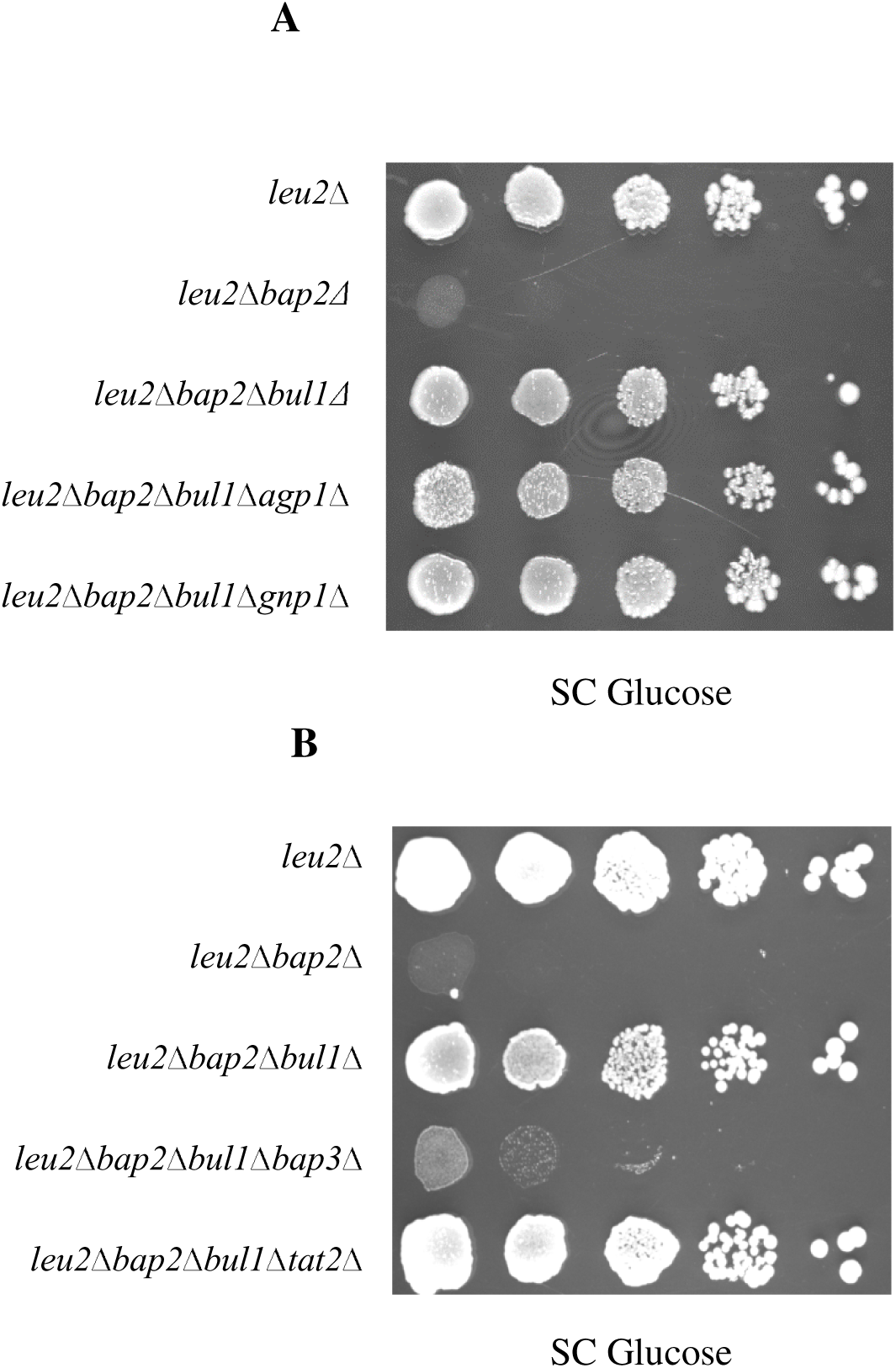
*BUL1* deletion suppresses the *leu2*Δ*bap2*Δ lethality in a *BAP3* dependent manner. (A) *leu2*Δ, *leu2*Δ*bap2*Δ, *leu2*Δ*bap2*Δ*bul1*Δ, *leu2*Δ*bap2*Δ*bul1*Δ*agp1*Δ and *leu2*Δ*bap2*Δ*bul1*Δ*gnp1*Δ strains grown in SC Glycerol + lactate broth were spotted on SC Glucose. Images were captured after incubating the petri plates for 2-3 days at 30°C. (B) *leu2*Δ, *leu2*Δ*bap2*Δ, *leu2*Δ*bap2*Δ*bul1*Δ, *leu2*Δ*bap2*Δ*bul1*Δ*bap3*Δ and *leu2*Δ*bap2*Δ*bul1*Δ*tat2*Δ strains grown in SC Glycerol + lactate broth were spotted on SC Glucose.

**Figure S2.**
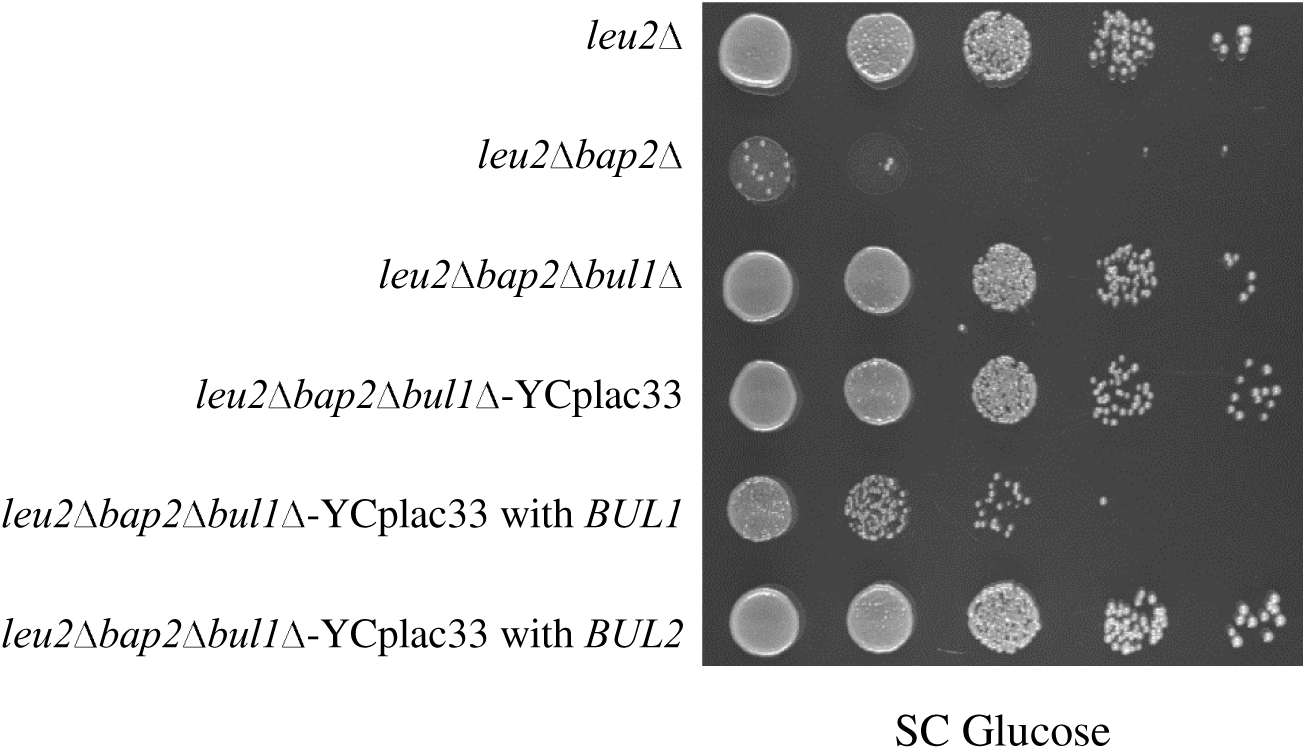
Complementation by *BUL1* not *BUL2* confers the lethality in *leu2*Δ*bap2*Δ*bul1*Δ strain. *leu2*Δ, *leu2*Δ*bap2*Δ, *leu2*Δ*bap2*Δ*bul1*Δ strains and *leu2*Δ*bap2*Δ*bul1*Δ transformant carrying YCplac33 or YCplac33 with *BUL1* or *BUL2* grown in SC Glycerol + lactate or S Ura^-^Glycerol + lactate broth was spotted for growth on SC Glucose. Images were captured after incubating the petri plates for 2-3 days at 30°C.

**Figure S3.**
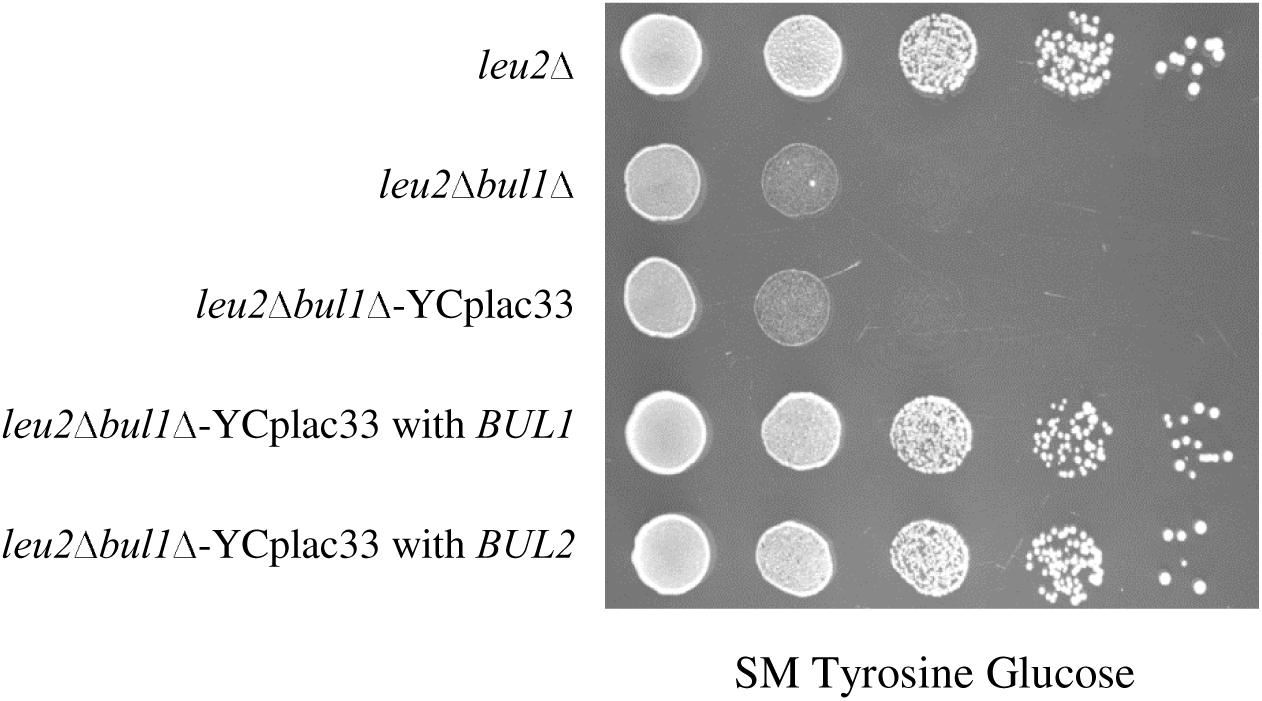
*BUL2* of W303-1A strain is functional. *leu2*Δ, *leu2*Δ*bul1*Δ strains and *leu2*Δ*bul1*Δ transformant carrying YCplac33 or YCplac33 with *BUL1*/*BUL2* grown in SC Glycerol + lactate or S Ura^-^ Glycerol + lactate broth was spotted for growth on SM Tyrosine Glucose. Images were captured after incubating the petri plates for 2-3 days at 37°C.

## Materials and methods

### Yeast strains and plasmids

All yeast strains are of BY4741 background (ST.1). They were either purchased from Euroscarf or constructed. Yeast transformation was performed by LiAc-PEG method. Yeast genomic DNA was isolated by glass bead method. Genetic manipulations like disruption and tagging were performed by PCR based homologous recombination method. These manipulations were again confirmed by PCR. Primers used for the above procedures are listed (ST.3).

E.coli strain XL1 or DH5α were used for plasmid cloning and propagation. Plasmid isolation was done by alkaline lysis method and *E.Coli* transformation was done by PEG method. Plasmids used in the study are listed (ST.2). Plasmid carrying *BUL1* was constructed by cloning of 4020 bp SalI-EcoRI fragment of *BUL1* gene (1000 bp upstream of start codon, 2931 bp CDS and 89 bp downsream of stop codon), into SalI-EcoRI site of YCplac33. *BUL1* was amplified from the genomic DNA of WT BY4741. Plasmid carrying *BUL2*, was constructed by cloning of a 4019 bp SalI-SacI fragment of *BUL2* (1000 bp upstream of start codon, 2763 bp CDS, 250 bp downsream of stop codon, 6 bp of primer region), amplified from genomic DNA of WT W303-1A strain, into the SalI-SacI site of YCplac33.

**Make –** Primers (Sigma, BLOT solutions), Restriction enzymes, polymerases and ligases (Thermo scientific, NEB), PCR/Gel purification kits (Qiagen).

### Media composition

**Synthetic Complete Glucose** or **Synthetic Complete Galactose (SC Glucose/SC Galactose)** – 0.17% Yeast nitrogen base, 0.5% ammonium sulfate, 0.05% amino acid mixture (ST4) and 2% glucose or galactose. **Synthetic Complete Glycerol (SC Glycerol)** – 0.17% Yeast nitrogen base, 0.5% ammonium sulfate, 0.05% amino acid mixture and 3% (V/V) glycerol. **Synthetic Complete Glycerol plus lactate (SC Glycerol + lactate)** – 0.17% Yeast nitrogen base, 0.5% ammonium sulfate, 0.05% amino acid mixture and 3% (V/V) glycerol and 2% (V/V) of 40%

lactate pH 5.7. **Synthetic Complete Ethanol (SC Ethanol)** – 0.17% Yeast nitrogen base, 0.5% ammonium sulfate, 0.05% amino acid mixture and 4% (V/V). SUra^-^Glycerol + lactate and SLeu^-^ Glycerol + lactate refers to SCGlycerol + lactate lacking uracil or leucine respectively. **Synthetic Minimal Tyrosine Glucose (SM Tyrosine Glucose)** – 0.17% Yeast nitrogen base, 2% glucose, 0.01% of auxotrophic supplements and 2mM Tyrosine. For solid media 2%agar was used. **YPD -** 2% glucose, 1% bacto peptone and 0.5% yeast extract.

### Spotting assay (Growth assay)

Cells were grown in SCGlycerol + lactate broth (SCGlycerol + lactate broth lacking uracil or leucine for transformants) to mid log phase, harvested and washed with sterile distilled water and cell density was normalized to OD_600_ 1.0. The normalized cell suspension was serially diluted to tenfold dilutions up to 10^-4^ dilution. 5µl from each dilution was spotted onto appropriate media and incubated at 30°C or 37°C for 2-3 days. Images of the petriplates were captured by UVITEC, CAMBRIDGE gel documentation system of GeNei.

### Protein extraction and immunoblotting

Protein was extracted by alkaline lysis method. Cells grown in appropriate medium to mid log phase were harvested at 4°C and lysed with 0.5ml of lysis solution (0.2M NaOH, 0.2% mercaptoethanol) for 10 min on ice. Trichloroacetic acid was added to a final concentration of 5%, and incubated for another 10 min on ice. Samples were centrifuged for 5 min at 13000 x g for 5 min. Pellets were resuspended in 35 µl of dissociation buffer (4% SDS, 0.1M Tris-HCl pH 6.8, 4mM EDTA, 20% glycerol, 2% 2-mercaptoethanol, 0.02% Bromophenol blue) and mixed with 15 µl of 1M Tris base, and heated at 37°C for 20 min.

Total cell free extracts were resolved by SDS-PAGE and transferred to 0.45 micron nitrocellulose membrane. Mouse bi-clonal anti-GFP antibodies were used to probe GFP tagged proteins and chased by secondary anti-mouse goat antibody conjugated with alkaline phosphatase. Rabbit anti-G6PD antibody was used to probe G6PD, and chased down by secondary anti-rabbit mouse antibody conjugated with alkaline phosphatase. Blots were developed and images were captured by UVITEC, CAMBRIDGE gel documentation system of GeNei and analyzed by UVITEC1D software.

**Make –** Mouse bi-clonal anti-GFP antibody (Roche Diagnostics, Mix of 13.1 and 7.1 monoclonal antibodies), Rabbit anti-G6PD antibody (Santacruz Biotechnology) and Secondary anti-rabbit mouse antibody (Santacruz Biotechnology). Nitrocellulose membrane (Amersham Biosciences). SDS-PAGE and blotting transfer apparatus (GeNei).

### Fluorescence microscopy

Strains were grown in appropriate media to mid log phase were harvested, washed in sterile distilled water and resuspended in the same (To avoid auto fluorescence due to nutrient broth). Cells were readily analyzed for GFP fluorescence at room temperature under Zeiss Axio-Observer Z1 inverted motorized and computer-controlled laser scanning fluorescence microscope (Carl Zeiss, LSM 780) equipped with iPlan-Apochromat 100X oil immersion objective 1.4 NA and PMT detector camera. Argon+ laser was used to excite the fluorophore with excitation wavelength of 488nm and emission wavelength was 526nm. Images were captured and later processed by Zen 2012 blue acquisition software from Zeiss.

### Leucine import assay

Strains were grown in SC Glycerol + lactate broth lacking leucine; sub-cultured in SC Glucose broth and grown up to mid log phase (OD_600_ 1-1.5). Cells were harvested, washed with SC Glucose broth twice and resuspended in the same to 0.4×10^7^cells/ml. To 200µl of this suspension added was 2.8KBq (500 picomoles) of C14-labelled leucine. The suspension contained 188µM of cold leucine and 2.5µM of C14-leucine. The cell suspensions were incubated in 30°C incubator cum shaker for 0 min, 60 min and 120 min time points. The C14-labelled leucine import was stopped by adding cold leucine of 100µl to a final leucine concentration of 12.6mM. The cells from the suspension were collected on a 0.45 micron nylon membrane (Amersham Biosciences) by suction. Cells were washed with ice cold water for 3 times and the membrane was dried for 10 minutes. The membrane with cells was counted for radioactive pulses for 1min/sample in Perkin Elmer Liquid scintillation analyzer. For each repetition biological replicates were taken and technical triplicates were taken for each strain for each time point. C14-leucine with specific activity of 5.5GBq/mmole (150mci/mmole) was purchased from BRIT, India.

## Supplementary tables

**ST1.**
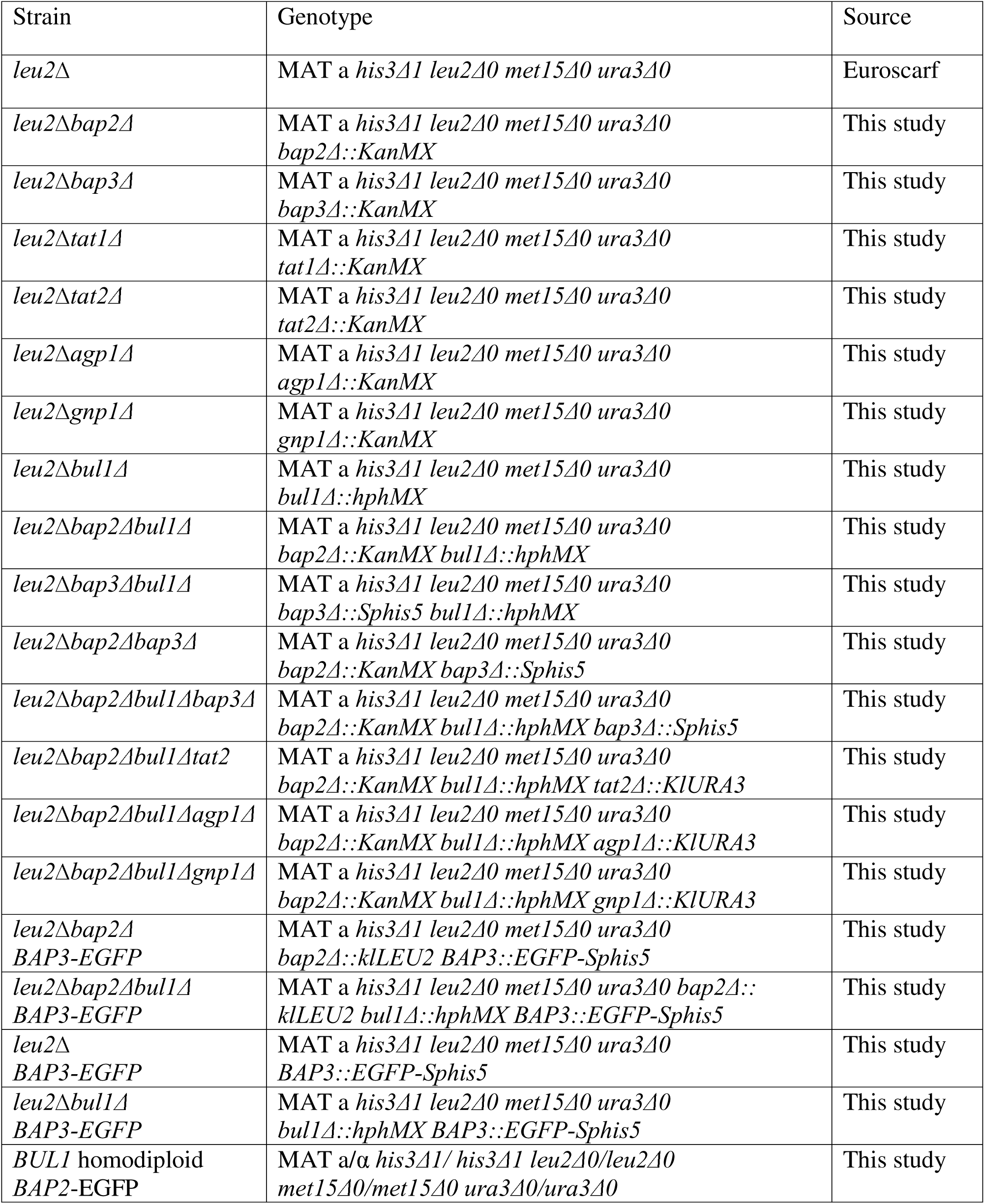
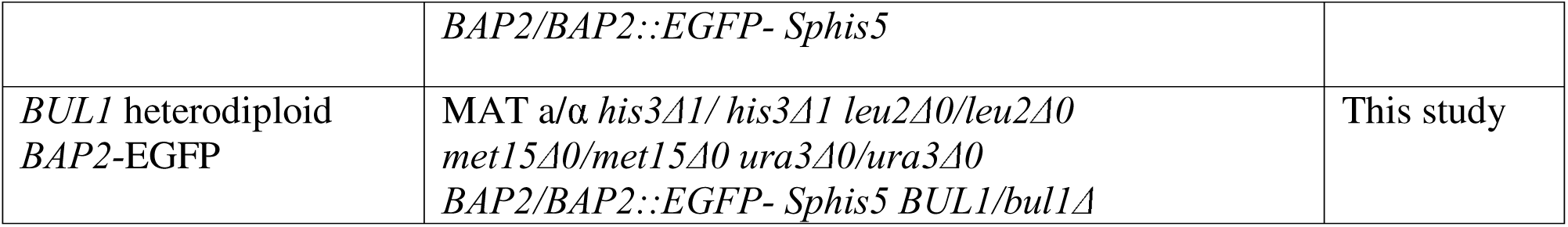
Yeast strains.

**ST2.** Plasmids.

ST2. Plasmids.
| Plasmid | Description | Source |
| --- | --- | --- |
| YCplac33 | Yeast centromeric shuttle vector with *AMP<sup>R</sup>* and *URA3* markers | Lab stock |
| YCplac33-*BUL1* | Yeast centromeric shuttle vector with *AMP<sup>R</sup>* and *URA3* markers carrying *BUL1* | This study |
| YCplac33-*BUL2* | Yeast centromeric shuttle vector with *AMP<sup>R</sup>* and *URA3* marker carrying *BUL2* | This study |
| YEplac181 | Yeast episomal shuttle vector with *AMP<sup>R</sup>* and *LEU2* markers | Lab stock |

**ST3.**
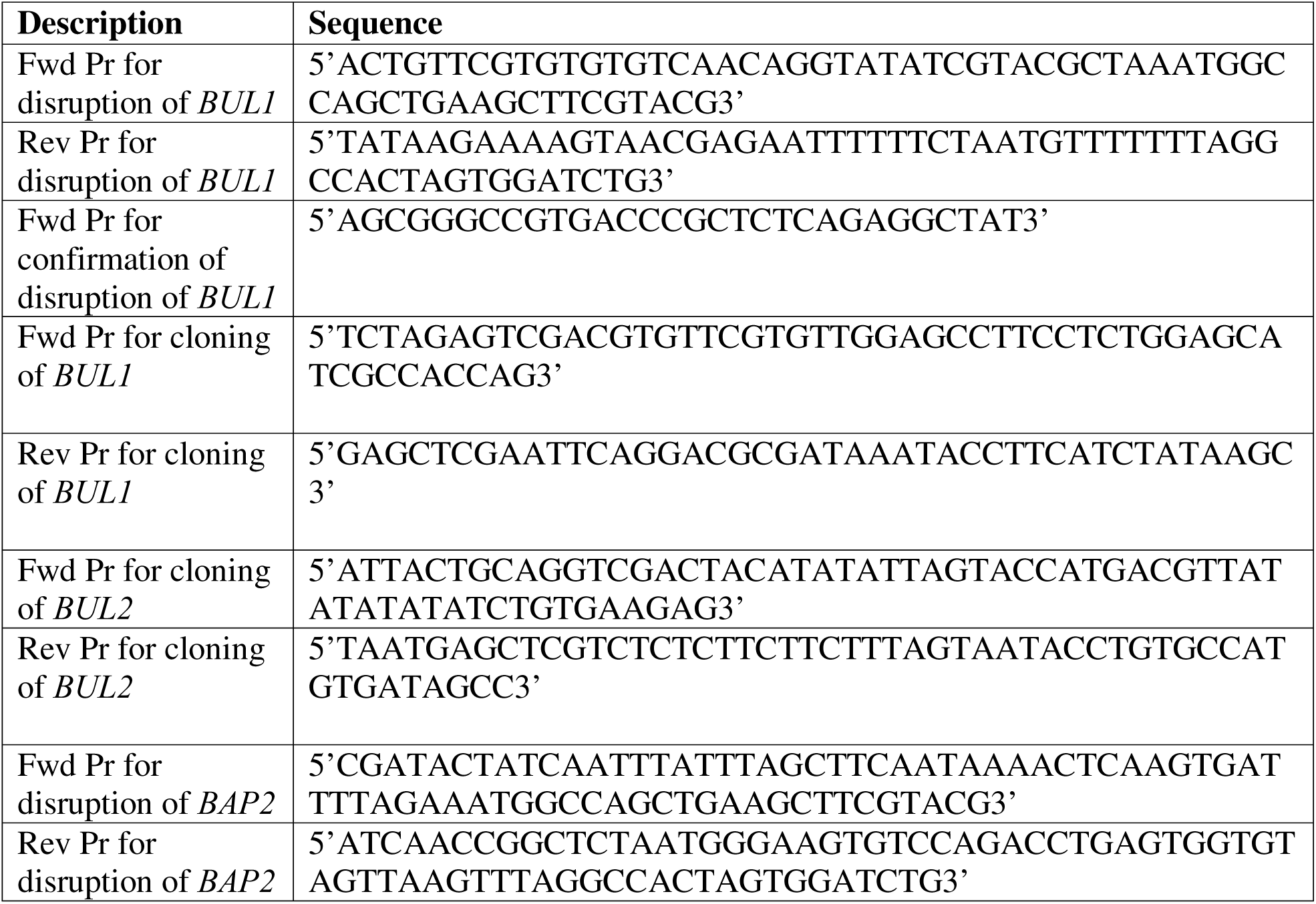
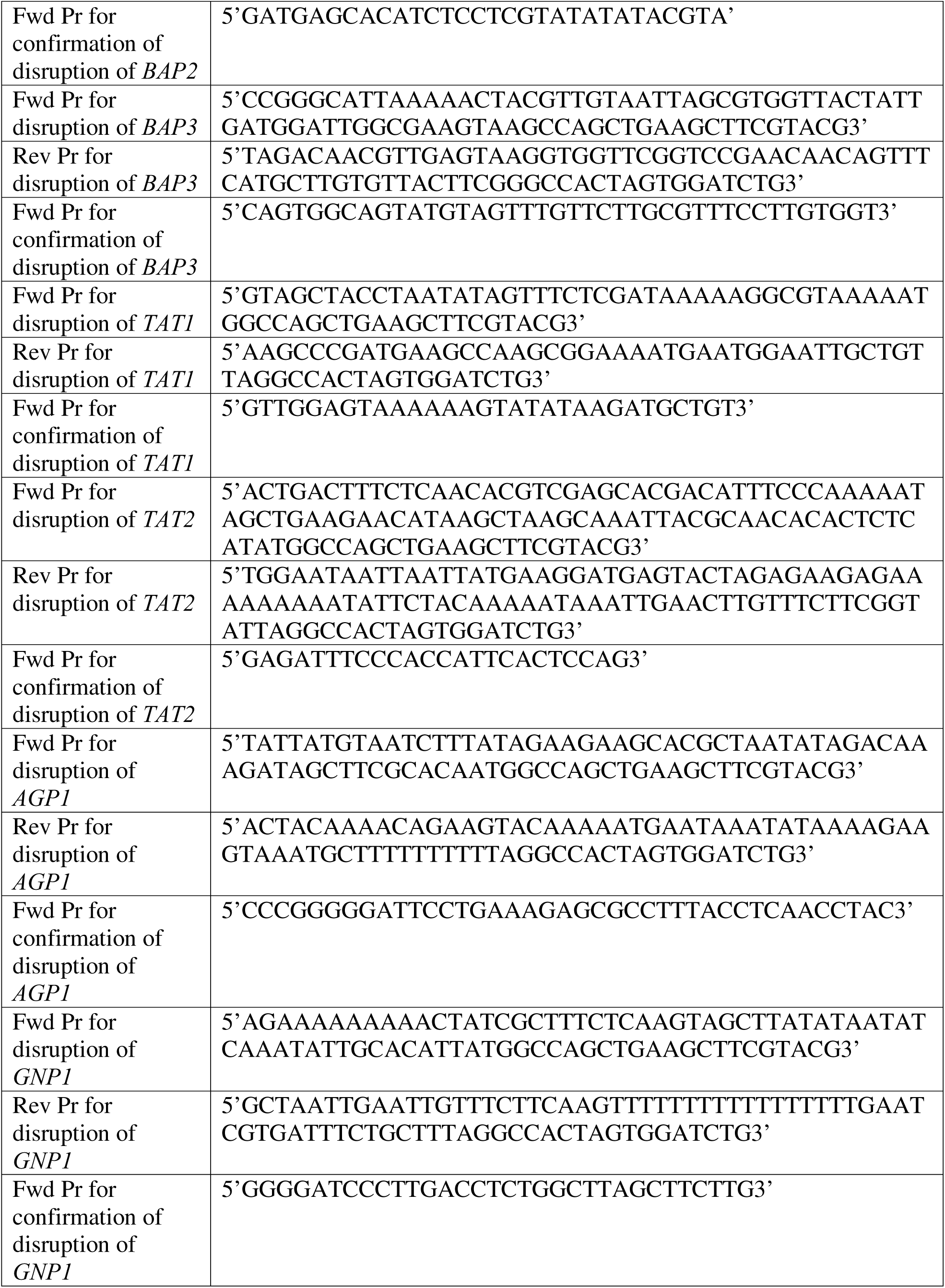
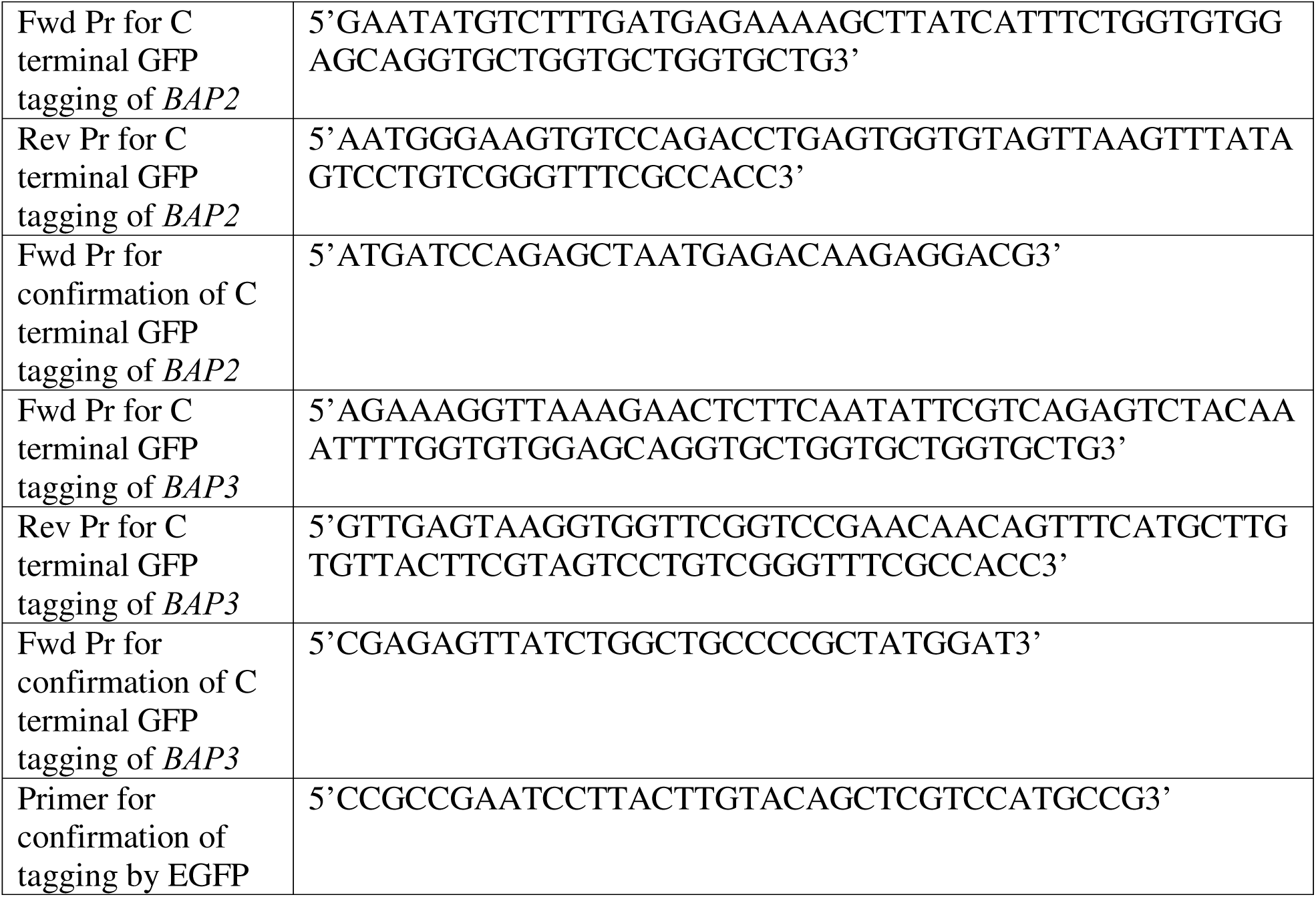
Primers.

## Acknowledgements

We are very grateful to Department of Science and Technology, INDIA, for funding the project (P13DST024).

## Conflict of interest

The authors declare no conflict of interest.

## Author contribution

MS and PJB conceived ideas and designed the experiments. MS performed the experiments. MS and PJB analyzed the results. MS and PJB wrote the manuscript.

## Data availability statement

Data supporting the findings of this study are available, upon request.

## Notes

### Competing Interest Statement

The authors have declared no competing interest.

## References

Abe, F., and H. Iida. 2003. Pressure induced differential regulation of the two tryptophan permeases Tat1 and Tat2 by ubiquitin ligase Rsp5 and its binding proteins, Bul1 and Bul2. Mol. Cell. Biol. 23:7566–7584.

Alvers, A.L., L.K. Fishwick, M.S. Wood, D. Hu, H.S. Chung, W.A. Dunn Jr and J.P. Aris. 2009. Autophagy and amino acid homeostasis are required for chronological longevity in *Saccharomyces cerevisiae*. Aging cell. 8(4):353–369.

Beck, T., A. Schmidt, and M.N. Hall. 1999. Starvation induces vacuolar targeting and degradation of the tryptophan permease in yeast. J Cell Biol. 146:1227–38.

Becuwe, M., N. Vieira, D. Lara, J. Gomes-Rezende, C. Soares-Cunha, M. Casal, R. Haguenauer-Tsapis, O. Vincent, S. Paiva, and Léon S. 2012. A molecular switch on an adaptor-like protein relays glucose signaling to transporter endocytosis. J. Cell Biol. 196:247–259.

Bianchi, F., J.S.V. Klooster, S.J. Ruiz, and B. Poolman. 2019. Regulation of Amino Acid Transport in *Saccharomyces cerevisiae*. Microbiol Mol Biol Rev. 4:e00024–19.

Blondel, M.O., J. Morvan, S. Dupré, D. Urban-Grimal, R. Haguenauer-Tsapis, and C. Volland. 2004. Direct sorting of the yeast uracil permease to the endosomal system is controlled by uracil binding and Rsp5p-dependent ubiquitylation. Mol Biol Cell. 15:883–95.

Bonfils, G., M. Jaquenoud, S. Bontron, C. Ostrowicz, C. Ungermann, and C. De Virgilio. 2012. Leucyl-tRNA synthetase controls TORC1 via the EGO complex. Mol Cell. 46:105–110.

Cohen, R., and D. Engelberg. 2007. Commonly used *Saccharomyces cerevisiae* strains (e.g. BY4741, W303) are growth sensitive on synthetic complete medium due to poor leucine import. FEMS Microbiol Lett. 273:239–243.

Crapeau, M., A. Merhi, and B. André. 2014. Stress conditions promote yeast Gap1 permease ubiquitylation and down-regulation via the adaptor-like Bul and Aly proteins. J. Biol. Chem. 289:22103–22116.

Didion, T., B. Regenberg, M.U. Jorgensen, M.C. Kielland-Brandt, and H.A. Andersen. 1998. The permease homologue Ssy1p controls the expression of amino acid and peptide transporter genes in *Saccharomyces cerevisiae*. Mol Microbiol. 27:643–50.

Echols, N., P. Harrison, S. Balasubramanian, N.M. Luscombe, P. Bertone, Z. Zhang, and M. Gerstein. 2002. Comprehensive analysis of amino acid and nucleotide composition in eukaryotic genomes, comparing genes and pseudogenes. Nucleic Acids Res. 30:2515–2523.

Esch, B.M., S. Limar, A. Bogdanowski, C. Gournas, T. More, C. Sundag, S. Walter, J. J Heinisch, C.S. Ejsing, B. André, and F. Fröhlich. 2020. Import of exogenous serine is important to maintain sphingolipid homeostasis in *Saccharomyces cerevisiae*. PLoS Genet. 16:e1008745.

Felice, M.R., I. De Domenico, L. Li, D.M. Ward, B. Bartok, G. Musci, and J. Kaplan. 2005. Post-transcriptional regulation of the yeast high affinity iron transport system. J Biol Chem. 280:22181–90.

Fernandez-Murray, J.P., M.H. Ngo, and C.R. McMaster. 2013. Choline transport activity regulates phosphatidylcholine synthesis through choline transporter Hnm1 stability. J Biol Chem. 50:36106–15.

Fujita, S., D. Sato, H. Kasai, M. Ohashi, S. Tsukue, Y. Takekoshi, K. Gomi, and T. Shintani. 2018. The C-terminal region of the yeast monocarboxylate transporter Jen1 acts as a glucose signal-responding degron recognized by the α-arrestin Rod1. J Biol Chem. 293:10926–10936.

Ghaemmaghami, S., W.K. Huh, K. Bower, R.W. Howson, A. Belle, N. Dephoure, E.K. O’Shea, and J.S. Weissman. 2003. Global analysis of protein expression in yeast. Nature. 425:737–741.

Gitan. R.S and Eide. D.J. 2000. Zinc-regulated ubiquitin conjugation signals endocytosis of the yeast ZRT1 zinc transporter. Biochem J. 346:329–36.

Grauslund, M., T. Didion, M.C. Kielland-Brandt, and H.A. Andersen. 1995. *BAP2*, a Gene Encoding a Permease for Branched-Chain Amino Acids in *Saccharomyces Cerevisiae*. Biochim Biophys Acta. 1269:275–80.

Han, J.M., S.J. Jeong, M.C. Park, G. Kim, N.H. Kwon, H.K. Kim, S.H. Ha, S.H. Ryu, and S. Kim. 2012. Leucyl tRNA synthetase is an intracellular leucine sensor for the mTORC1 signaling pathway. Cell. 149:410–424.

Hatakeyama, R., M. Kamiya, T. Takahara, and T. Maeda. 2010. Endocytosis of the aspartic acid/glutamic acid transporter Dip5 is triggered by substrate-dependent recruitment of the Rsp5 ubiquitin ligase via the adaptor-like protein Aly2. Mol. Cell. Biol. 30:5598–5607.

Hettema, E.H., J. Valdez-Taubas, and H.R. Pelham. 2004. Bsd2 binds the ubiquitin ligase Rsp5 and mediates the ubiquitination of transmembrane proteins. EMBO J. 23:1279–88.

Horak. J. and Wolf. D.H. 1997. Catabolite inactivation of the galactose transporter in the yeast *Saccharomyces cerevisiae*: ubiquitination, endocytosis, and degradation in the vacuole. J Bacteriol. 179:1541–9.

Hovsepian, J., Q. Defenouillere, V. Albanese, L. Vachova, C. Garcia, Z. Palkova, and S. Leon. 2017. Multilevel regulation of an alpha-adaptor by glucose depletion controls hexose transporter endocytosis. J. Cell Biol. 216:1811–1831.

Hovsepian, J., V. Albanese, M. Becuwe, V. Ivashov, D. Teis, and S. Leon. 2018. The yeast adaptor-related protein Bul1 is a novel actor of glucose-induced endocytosis. Mol. Biol. Cell. 29:1012–1020.

Leon, S., Z. Erpapazoglou, and R. Haguenauer-Tsapis. 2008. Ear1p and Ssh4p are new adaptors of the ubiquitin ligase Rsp5p for cargo ubiquitylation and sorting at multivesicular bodies. Mol Biol Cell. 19:2379–88.

Liu, X.F., and V.C. Culotta. 1999. Post-translation control of Nramp metal transport in yeast. Role of metal ions and the BSD2 gene. J Biol Chem. 274:4863–8.

Kahlhofer, J., S. Leon, D. Teis, and O. Schmidt. 2021. The α-adaptor family of ubiquitin ligase adaptors links metabolism with selective endocytosis. Biol Cell. 4:183–219.

Kozu, F., K. Shirahama-Noda, Y. Araki, S. Kira, H. Niwa, and T. Noda. 2021. Isoflurane induces Art2-Rsp5-dependent endocytosis of Bap2 in yeast. FEBS Open Bio. 11:3090–3100.

Krampe, S., O. Stamm, C.P Hollenberg, and E. Boles. 1998. Catabolite inactivation of the high-affinity hexose transporters Hxt6 and Hxt7 of *Saccharomyces cerevisiae* occurs in the vacuole after internalization by endocytosis. FEBS Lett. 441:343–7.

Kwan, E.X., E. Foss, L. Kruglyak, and A. Bedalov. 2011. Natural Polymorphism in *BUL2* Links Cellular Amino Acid Availability With Chronological Aging and Telomere Maintenance in Yeast. Plos Genet. 7(8):e1002250.

Lai, K., C.P. Bolognese, S. Swift, and P. McGraw. 1995. Regulation of inositol transport in *Saccharomyces cerevisiae* involves inositol-induced changes in permease stability and endocytic degradation in the vacuole. J Biol Chem. 270:2525–34.

Lin, C.H., J.A. MacGurn, T. Chu, C.J Stefan, and S.D. Emr. 2008. Adaptor related ubiquitin ligase adaptors regulate endocytosis and protein turnover at the cell surface. Cell. 135:714–725.

Liu, J., A. Sitaram, and C.G. Burd. 2007. Regulation of copper-dependent endocytosis and vacuolar degradation of the yeast copper transporter, Ctr1p, by the Rsp5 ubiquitin ligase. Traffic. 8:1375–84.

Megarioti, A.H., C. Primo, G.C. Kapetanakis, A. Athanasopoulos, V. Sophianopoulou, B. Andre, and C. Gournas. 2021. The Bul1/2 Alpha-Arrestins Promote Ubiquitylation and Endocytosis of the Can1 Permease upon Cycloheximide Induced TORC1 Hyperactivation. Int J Mol Sci. 19:10208.

Merhi, A., and B. Andre. 2012. Internal amino acids promote Gap1 permease ubiquitylation via TORC1/Npr1/14-3-3-dependent control of the Bul adaptor-like adaptors. Mol. Cell. Biol. 32: 4510–4522.

Nigavekar. S.S. and Cannon. J.F. 2002. Characterization of genes that are synthetically lethal with ade3 or leu2 in *Saccharomyces cerevisiae*. Yeast. 2:115–22.

Nikko, E., J.A. Sullivan, and H.R. Pelham. 2008. Adaptor-like proteins mediate ubiquitination and endocytosis of the yeast metal transporter Smf1. EMBO Rep. 9:1216–1221.

Nikko, E., and H.R. Pelham. 2009. Adaptor-mediated endocytosis of yeast plasma membrane transporters. Traffic. 10:1856–1867.

O’Donnell, A.F. 2012. The Running of the Buls: Control of Permease Trafficking by α-Adaptors Bul1 and Bul2. Mol Cell Biol. 32:4506–4509.

O’Donnell, A.F., and M.C. Schmidt. 2019. AMPK-Mediated Regulation of Alpha-Adaptors and Protein Trafficking. Int J Mol Sci. 20(3):515.

Paiva, S., N. Vieira, I. Nondier, R. Haguenauer-Tsapis, M. Casal, and D. Urban-Grimal. 2009. Glucose-induced ubiquitylation and endocytosis of the yeast Jen1 transporter: role of lysine 63-linked ubiquitin chains. J Biol Chem. 284:19228–36.

Peter, G.J., L. Düring, and A. Ahmed. 2006. Carbon Catabolite Repression Regulates Amino Acid Permeases in *Saccharomyces cerevisiae* via the TOR Signaling Pathway. J Biol Chem. 281(9):5546–52.

Regenberg, B., L. During-Olsen, M.C. Kielland-Brandt, and S. Holmberg. 1999. Substrate specificity and gene expression of the amino-acid permeases in *Saccharomyces cerevisiae*. Curr Genet. 36:317–28.

Robinson, B.P., S. Hawbaker, A. Chiang, E.M. Jordahl, S. Anaokar, A. Nikiforov, R.W. Bowman, P. Ziegler, C.K. McAtee, J. Patton-Vogt, and A.F. O’Donnell. 2022. Alpha-arrestins Aly1/Art6 and Aly2/Art3 regulate trafficking of the glycerophosphoinositol transporter Git1 and impact phospholipid homeostasis. Biol Cell. 114:3–31.

Sardana. R. and Emr. S.D. 2021. Membrane Protein Quality Control Mechanisms in the Endo-Lysosome System. Trends Cell Biol. 31:269–283.

Savocco, J., S. Nootens, W. Afokpa, M. Bausart, X. Chen, J. Villers, H. Renard, M. Prévost, R. Wattiez, and P. Morsomme. 2019. Yeast α-Adaptor Art2 Is the Key Regulator of Ubiquitylation- Dependent Endocytosis of Plasma Membrane Vitamin B1 Transporters. Plos Biol. 17(10):e3000512.

Segev. N and Hay. N. 2012. Hijacking leucyl-tRNA synthetase to amino acid-dependent regulation of TORC1. Mol Cell. 46:4–6.

Stimpson, H.E., M.J. Lewis, and H.R. Pelham. 2006. Transferrin receptor-like proteins control the degradation of a yeast metal transporter. EMBO J. 25:662–72.

Suzuki, A., T. Mochizuki, S. Uemura, T. Hiraki, and F. Abe. 2013. Pressure-induced endocytic degradation of the *Saccharomyces cerevisiae* low-affinity tryptophan permease Tat1 is mediated by Rsp5 ubiquitin ligase and functionally redundant PPxY motif proteins. Eukaryot. Cell. 12:990–997.

Tanahashi, R., T. Matsushita, A. Nishimura, and H. Takagi. 2021. Downregulation of the broad-specificity amino acid permease Agp1 mediated by the ubiquitin ligase Rsp5 and the adaptor-like protein Bul1 in yeast. Biosci Biotechnol Biochem. 85:1266–1274.

Talaia, G., C. Gournas, E. Saliba, C. Barata-Antunes, M. Casal, B. Andre, G. Diallinas, and S. Paiva. 2017. The alpha-Adaptor Bul1p Mediates Lactate Transporter Endocytosis in Response to Alkalinization and Distinct Physiological Signals. J. Mol. Biol. 429:3678–3695.

Umebayashi. K and Nakano. A. 2003. Ergosterol is required for targeting of tryptophan permease to the yeast plasma membrane. J. Cell Biol. 161:1117–1131.

Usami, Y., S. Uemura, T. Mochizuki, A. Morita, F. Shishido, J. Inokuchi, and F. Abe. 2014. Functional mapping and implications of substrate specificity of the yeast high-affinity leucine permease Bap2. Biochim Biophys Acta. 7:1719–29.

Yashiroda, H., D. Kaida, A. Toh-e, and Y. Kikuchi. 1998. The PY-motif of Bul1 protein is essential for growth of *Saccharomyces cerevisiae* under various stress conditions. Gene. 225:39–46.

Zbieralski. K. and Wawrzycka. D. 2022. α-Arrestins and Their Functions: From Yeast to Human Health. Int J Mol Sci. 23:4988.

Zhang, W., G. Du, J. Zhou, and Jian Chen. 2018. Regulation of Sensing, Transportation, and Catabolism of Nitrogen Sources in *Saccharomyces cerevisiae*. Microbiol Mol Biol Rev. 82(1).

Zhao, Y., J.A. Macgurn, M. Liu, and S.D. Emr. 2013. The ART-Rsp5 ubiquitin ligase network comprises a plasma membrane quality control system that protects yeast cells from proteotoxic stress. eLife. 2:e00459.

